# Extraordinary preservation of gene collinearity over three hundred million years revealed in homosporous lycophytes

**DOI:** 10.1101/2023.07.24.548637

**Authors:** Cheng Li, David Wickell, Li-Yaung Kuo, Xueqing Chen, Bao Nie, Xuezhu Liao, Dan Peng, Jiaojiao Ji, Jerry Jenkins, Mellissa Williams, Shengqiang Shu, Chris Plott, Kerrie Barry, Shanmugam Rajasekar, Jane Grimwood, Xiaoxu Han, Shichao Sun, Zhuangwei Hou, Weijun He, Guanhua Dai, Cheng Sun, Jeremy Schmutz, James H. Leebens-Mack, Fay-Wei Li, Li Wang

## Abstract

Homosporous lycophytes (Lycopodiaceae) are a deeply diverged lineage in the plant tree of life, having split from heterosporous lycophytes (*Selaginella* and *Isoetes*) ∼400 million years ago (MYA). Compared to the heterosporous lineage, Lycopodiaceae has markedly larger genome sizes and remains the last major plant clade for which no genomic data has been available. Here, we present chromosomal genome assemblies for two homosporous lycophyte species, the allotetraploid *Huperzia asiatica* and the diploid *Diphasiastrum complanatum*. Remarkably, despite that the two species diverged ∼350 MYA, around 30% of the genes are still in syntenic blocks. Furthermore, both genomes had undergone independent whole genome duplications and the resulting intra-genomic syntenies have likewise been preserved relatively well. Such slow genome evolution over deep time is in stark contrast to heterosporous lycophytes and is correlated with a decelerated rate of nucleotide substitution. Together, the genomes of *H. asiatica* and *D. complanatum* not only fill a crucial gap in the plant genomic landscape, but also uncover a possibly unique genomic contrast between homosporous and heterosporous species.

## Introduction

Lycophytes occupy an important phylogenetic position sister to the euphyllophytes (ferns+seed plants), and have a rich evolutionary history with fossil records dating back to the late Devonian^1^. They have independently evolved many complex traits alongside ferns and seed plants including photosynthetic leaves and heterospory. As such, lycophytes represent a vital resource for understanding the early evolution of vascular plants on land^2^.

Extant lycophytes comprise around 1,330 species in three deeply diverged families, Selaginellaceae, Isoetaceae, and Lycopodiaceae^3^ (Fig. 1a). Like ferns, lycophytes also include both homosporous and heterosporous members, with Selaginellaceae and Isoetaceae being heterosporous, and Lycopodiaceae being entirely homosporous^4^. Homosporous plants produce only one type of spore, which develops into a bisexual gametophyte that produces both sperm and egg cells. On the other hand, heterosporous plants produce two types of spores (micro- and megaspore) that develop into separate male and female gametophytes. Homosporous lycophytes tend to have larger genome sizes and higher chromosome numbers than their heterosporous counterparts^5^. Whole genome duplication (WGD) has long been recognized as a key driver of genome size evolution and species diversification, and is relatively frequent in flowering plants^6–9^. Interestingly in lycophytes, the history of WGDs varies substantially across the major lineages. No ancient WGDs have been detected in Selaginellaceae^10–12^, and only one round of WGD has been conclusively demonstrated in Isoetaceae^13^. Conversely, several independent ancient WGDs have been postulated in Lycopodiaceae based on transcriptomic data^9^ but have yet to be verified with whole genome data. A thorough characterization of WGDs in Lycopodiaceae, especially regarding genome evolution post duplication, is needed to clarify what underlies the genome size disparity between heterosporous and homosporous lycophytes.

**Fig. 1.**
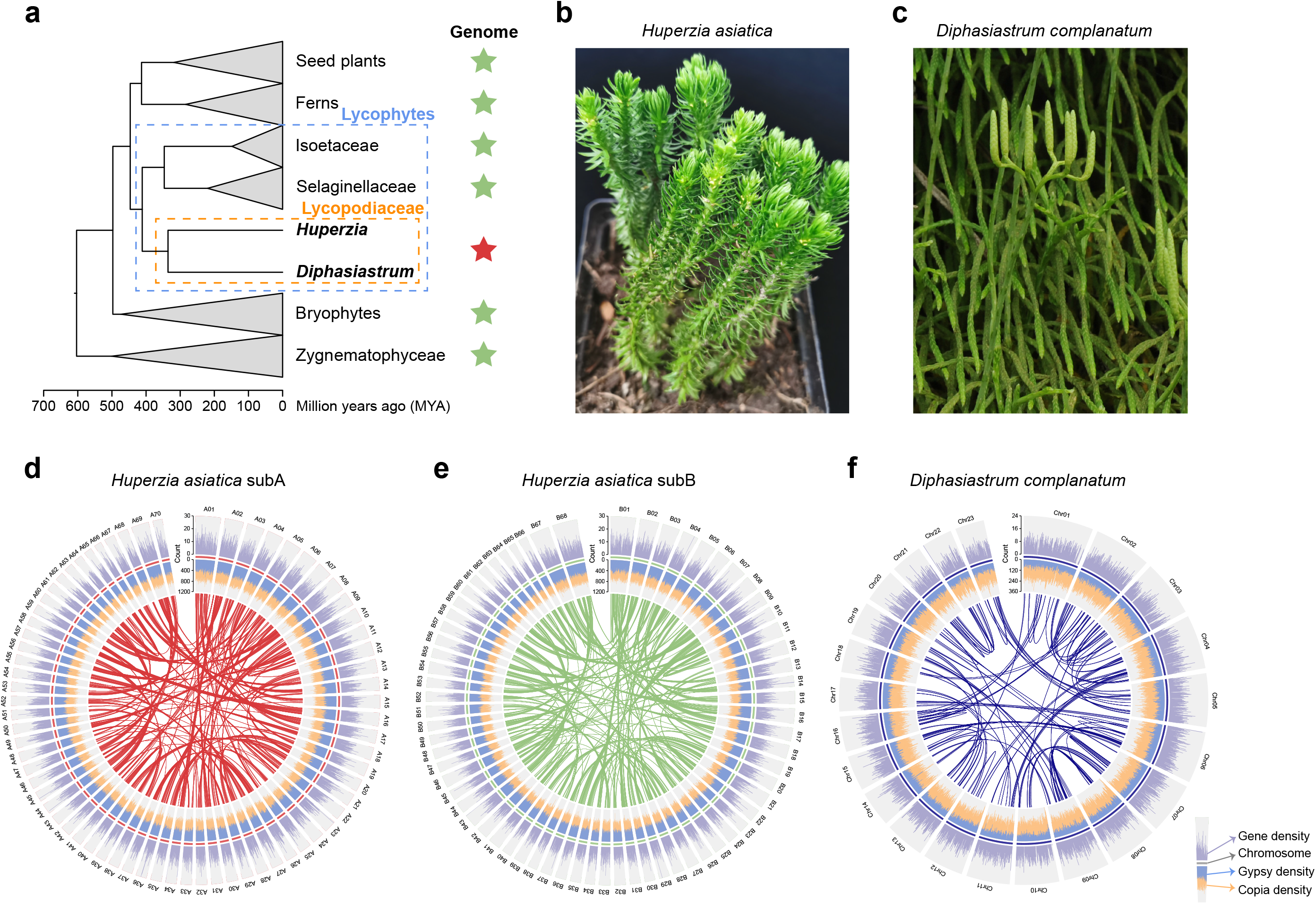
The genomes of *Huperzia asiatica* and *Diphasiastrum complanatum*. **a,** Homosporous lycophytes (Lycopodiaceae), which diverged from the closest extant group over 400 MYA, are the last major lineage for which no genome had been available. A green star indicates that at least one genome from this lineage has been sequenced, while a red star indicates that no genome has been sequenced. **b,c,** Whole genomes were assembled for *H. asiatica* (**b**) and *D. complanatum* (**c**). **d,e,f,** Genome features of *H. asiatica* and *D. complanatum*. The densities of genes and *gypsy* and *copia* LTRs for *H. asiatica* and *D. complanatum* were calculated with 1 Mb and 500 Kb non-overlapped sliding windows, respectively. Intra-genomic syntenies are displayed with connecting lines for subgenomes of *H. asiatica* and genome of *D. complanatum*. Image of *D. complanatum* courtesy of P.-F. Lu.

To date, the published lycophyte genomes have mainly focused on heterosporous families Selaginellaceae and Isoetaceae and none of them at the chromosomal level^10–14^. Given the deep divergences among these three lycophyte lineages (Fig. 1a), the lack of relatively complete genome assemblies has left a deep gap in our knowledge of lycophyte genomics and hindered the inferences of vascular plant evolution. Here we generated chromosome-level genome assemblies of the allotetraploid *Huperzia asiatica* and the diploid *Diphasiastrum complanatum*, belonging to Huperzioideae and Lycopodioideae subfamilies, respectively (Fig. 1b,c). We found a remarkably preserved synteny between these two genomes, despite the fact that the two species diverged over 350 million years ago (MYA) and have undergone multiple rounds of WGDs. Further, we found little bias in genome fractionation and homoeologous gene expression in *H. asiatica* after the most recent allotetraploidization event. Such slow genome evolution appears to correlate with slower rates of nucleotide substitutions compared to other lycophyte lineages. Our research fills a long-standing gap in land plant evolution and sheds light on the evolutionary history of early vascular plants tracing back hundreds of millions of years.

## Results and discussion

### Genome assembly and annotation

Based on k-mer and flow cytometry analyses, the sizes of the allotetraploid (*2n* = 4X = 276) *H. asiatica* and the diploid (*2n* = 2X = 46) *D. complanatum* genomes were estimated to be around 7.80 Gb and 1.60 Gb, respectively (Supplementary Fig. 1). Using a combination of PacBio CLR long reads, Illumina short reads, and Hi-C technology, we obtained chromosome-level assemblies for both genomes (Supplementary Fig. 2 and Supplementary Table 1). For *H. asiatica*, the genome was assembled into 138 pseudochromosomes and 2,191 unanchored scaffolds, with a total length of 7.94 Gb and N50 of 57.02 Mb. Using subgenome-specific k-mers, we further partitioned the allotetraploid *Huperzia* genome into A and B subgenomes (hereafter referred to as ‘subA’ and ‘subB’) (Supplementary Figs. 3-5). While we detected no evidence of recent large-scale homoeologous exchange between *H. asiatica* subgenomes based on k-mer sequences (Supplementary Fig. 4b), we could not exclude the possibility of ancient exchange shortly after allopolyploidization. The details on subgenome phasing are described in the Supplementary Text. For *D. complanatum*, the assembly consisted of 23 pseudochromosomes and 5,153 unanchored scaffolds, with a total length 1.74 Gb and N50 of 59.47 Mb. The mapping rates of genomic Illumina reads were over 98.8% and 95.0% against the *H. asiatica* and *D. complanatum* assemblies (Supplementary Table 2), respectively, indicating highly complete assemblies.

Compared to the published heterosporous lycophytes (*Isoetes taiwanensis*^13^, *Selaginella moellendorffii*^10^, *S. lepidophylla*^12^, and *S. tamariscina*^11^), our homosporous genomes comprised larger proportions of transposable elements (TEs) (81.86% for *H. asiatica* and 65.97% for *D. complanatum*), particularly long terminal repeats retrotransposons (LTR-RTs) (60.55% for *H. asiatica* and 53.68% for *D. complanatum*) (Supplementary Table 3). A total of 82,725 and 31,430 high confidence protein-coding genes were annotated for *H. asiatica* and *D. complanatum*, respectively (Supplementary Table 1). Both proteomes had high completeness scores from Benchmarking Universal Single-Copy Ortholog (BUSCO) (90.4% for *H. asiatica*; 97.5% for *D. complanatum*), indicating high annotation quality (Supplementary Table 4). We found that *H. asiatica* and *D. complanatum* have 37.9 and 16.1-fold longer introns than the heterosporous lycophytes, respectively (Supplementary Table 5), which is consistent with the positive correlation between intron length and genome size documented in other plant lineages^15–17^. Compared to heterosporous lycophytes, the introns in homosporous lycophytes contain higher numbers of LTR (mean number 7.8-25.1 vs. 1.6-3.1) as well as larger total span (mean length 3.4-11.1 Kb vs. 0.5-1.9 Kb) (Supplementary Table 6), suggesting LTR insertion contributed to the intron length difference between homosporous and heterosporous genomes.

The first chromosome-level assembly and annotation of lycophyte genomes allowed us to examine the distribution of repeats and genes in this lineage. In angiosperms, repeats and genes are unevenly distributed, with gene density increasing from centromere to telomere. However, we observe a different scenario in our genomes where the distribution of both appears to be homogeneous across the chromosomes (Fig. 1d-f). A similar distribution was also reported in chromosome-level assemblies of bryophytes^18,19^ and ferns^20^. Together, these results provide evidence that the genomic organization in seed-free plants might be quite distinct from seed plants.

### Origin and evolution of the allotetraploid *H. asiatica*

To determine the parentage and timeline of subgenome divergence in *H. asiatica*, we reconstructed phylogenetic trees using genome and transcriptome data of *Huperzia* spp. and outgroup species (Fig. 2a). We found that *H. asiatica* subA and subB were clustered into two groups, which were named A-genome clade and B-genome clade, respectively. The divergence time between A- and B-genome clades was estimated to be ∼35.4 MYA. *H. asiatica* subA and subB diverged from their closely related diploid species, *H. miyoshiana* and *H. lucidula*, ∼17.8 MYA and ∼19.9 MYA, respectively. In addition, the chloroplast phylogeny, which most likely tracks the maternal inheritance, further resolved that the A-genome clade is probably the maternal donor of *H. asiatica* (Fig. 2a).

**Fig. 2.**
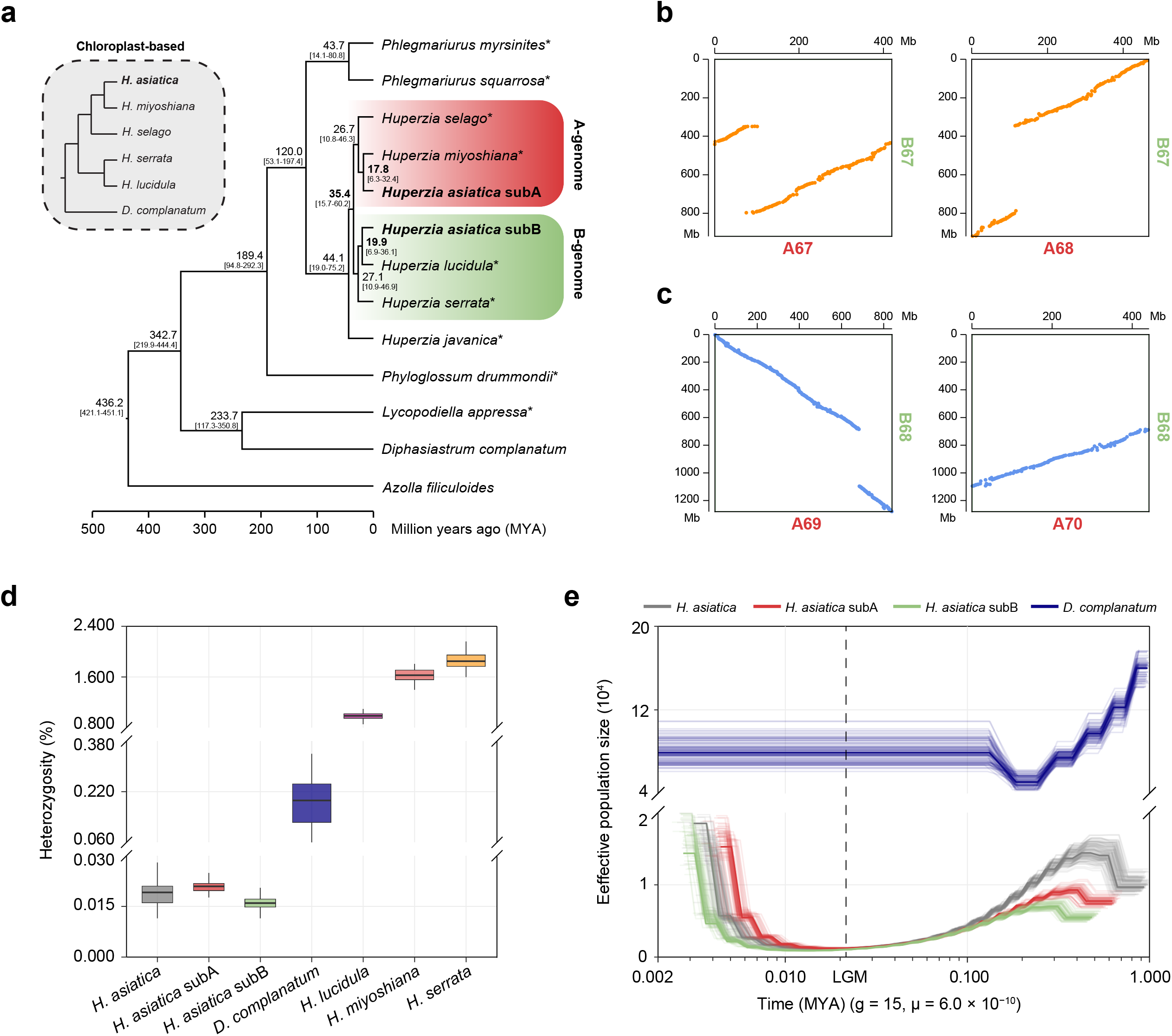
Origin and evolution of the allotetraploid *Huperzia asiatica*. **a,** Species phylogeny of Lycopodiaceae inferred from nuclear 1,288 gene families. The divergence times and the 95% confidence intervals of species or clades are labeled. Transcriptome-derived data are marked by an asterisk. The tree at upper left shows a plastome-based phylogeny from five *Huperzia* species and *Diphasiastrum complanatum*. **b,** Synteny maps show that the chromosome fusion or fission event involved in B67 and its subA homoeologs, A67 and A68, in allotetraploid *Huperzia* genome. **c,** Synteny maps show that the chromosome fusion or fission event involved in B68 and its subA homoeologs, A69 and A70, in allotetraploid *Huperzia* genome. **d,** Comparison of genome-wide heterozygosity estimated in four *Huperzia* species and *D. complanatum*. **e,** Demographic trajectory of *H. asiatica* and *D. complanatum*. Each curve represents one species, with thin lines depicting bootstraps. LGM, last glacial maximum.

Syntenic analysis of the two subgenomes of *H. asiatica* revealed 21,205 gene pairs in 649 collinear blocks (Supplementary Fig. 6) suggesting limited chromosomal rearrangement since divergence of the progenitor lineage genomes 35.4 MYA. Notably, only two large-scale chromosomal rearrangements (either chromosome fusion or fission events) were observed (Fig. 2b,c), which was also confirmed by Hi-C (Supplementary Fig. 2b). One rearrangement involved B67 and its homoeologs, A67 and A68, and the other one involved B68 and its homoeologs, A69 and A70. Interestingly, we found that the average expression levels of genes within the rearranged chromosomes were significantly higher than the genomic background (Supplementary Fig. 7a), which was not observed for the genes on the other chromosomes (Supplementary Fig. 7b). Gene Ontology (GO) enrichment analysis uncovered that the genes residing in these rearranged chromosomes were significantly enriched in functions associated with diversification of specialized metabolism, such as terpenoid and tocopherol biosynthesis (Supplementary Fig. 7c).

We observed that *H. asiatica* exhibited significantly lower genome-wide heterozygosity than other *Huperzia* species as well as *D. complanatum* (Fig. 2d; Wilcoxon rank-sum test *P*-value < 0.01; n = chromosome numbers). As low heterozygosity might be a consequence of long-term population bottlenecks^21,22^, we modeled changes in effective population size (N_e_) using the Pairwise Sequentially Markovian Coalescent method^23^. Overall, while both *H. asiatica* and *D. complanatum* had experienced a severe genetic bottleneck, the N_e_ of *H. asiatica* was significantly smaller (Fig. 2e). The ancestral N_e_ of *H. asiatica* was 1.5 x 10^4^ at ∼0.4 MYA and continuously declined until the end of the last glacial maximum (LGM; ∼20,000 years ago). Although N_e_ of *H. asiatica* underwent a rapid recovery from the LGM bottleneck, it remained at a low level compared to *D. complanatum*. The extremely small N_e_ and limited geographical distribution for an extended period echoed the endangered status of *H. asiatica*^24^.

### Limited subgenome dominance in *H. asiatica*

Studies in flowering plants have shown that following allopolyploidization, one of the subgenomes often rose to dominance, which can be in the form of preferential retention of homoeologs or elevated gene expression levels^25–28^. However, little is known about these processes outside of flowering plants. To compare the preferential gene retention after allopolyploidization, we characterized gene presence/absence variants (PAVs) between *H. asiatica* subgenomes. We found 3,742 (9.9% of the annotated genes in subA) and 4,570 (11.4% of the annotated genes in subB) orphan genes (i.e. lacking homoeologs) are present only in *H. asiatica* subA and subB, respectively. The average number of orphan genes per chromosome is 29.8 for subA and 31.0 for subB, and there is no significant difference between the subgenomes (Supplementary Fig. 8; Wilcoxon rank-sum test *P*-value = 0.52; n = chromosome numbers), suggesting a pattern of non-preferential retention of homoeologs. To quantify subgenome expression bias, we compared gene expression levels in 18,264 homoeologous gene pairs identified through synteny mapping (the light gray links in Supplementary Fig. 6 and Supplementary Table 8). Generally, we found a weak expression divergence between homoeologs with average log_2_(subB/subA) expression ratios ranging from 0.02 to 0.07 in different tissue types (Fig. 3a). Gene pairs exhibiting homoeologous expression bias (HEB) were identified with cutoffs ‘*P*-value < 0.05 and | log_2_FoldChange | > 1’. More than 70.0% of the 18,264 homoeolog pairs exhibited a non-biased expression pattern (Supplementary Fig. 9). For pairs with significant HEB, the more highly expressed homoeolog was equally distributed between subA and subB (Fig. 3b and Supplementary Fig. 10). Overall, the two subgenomes exhibit a relatively balanced pattern of homoeolog gene expression. Homoeologous exchange has been hypothesized to mask the influence of genome dominance on gene expression^29^, but here we find little evidence of it (Supplementary Fig. 4b).

**Fig. 3.**
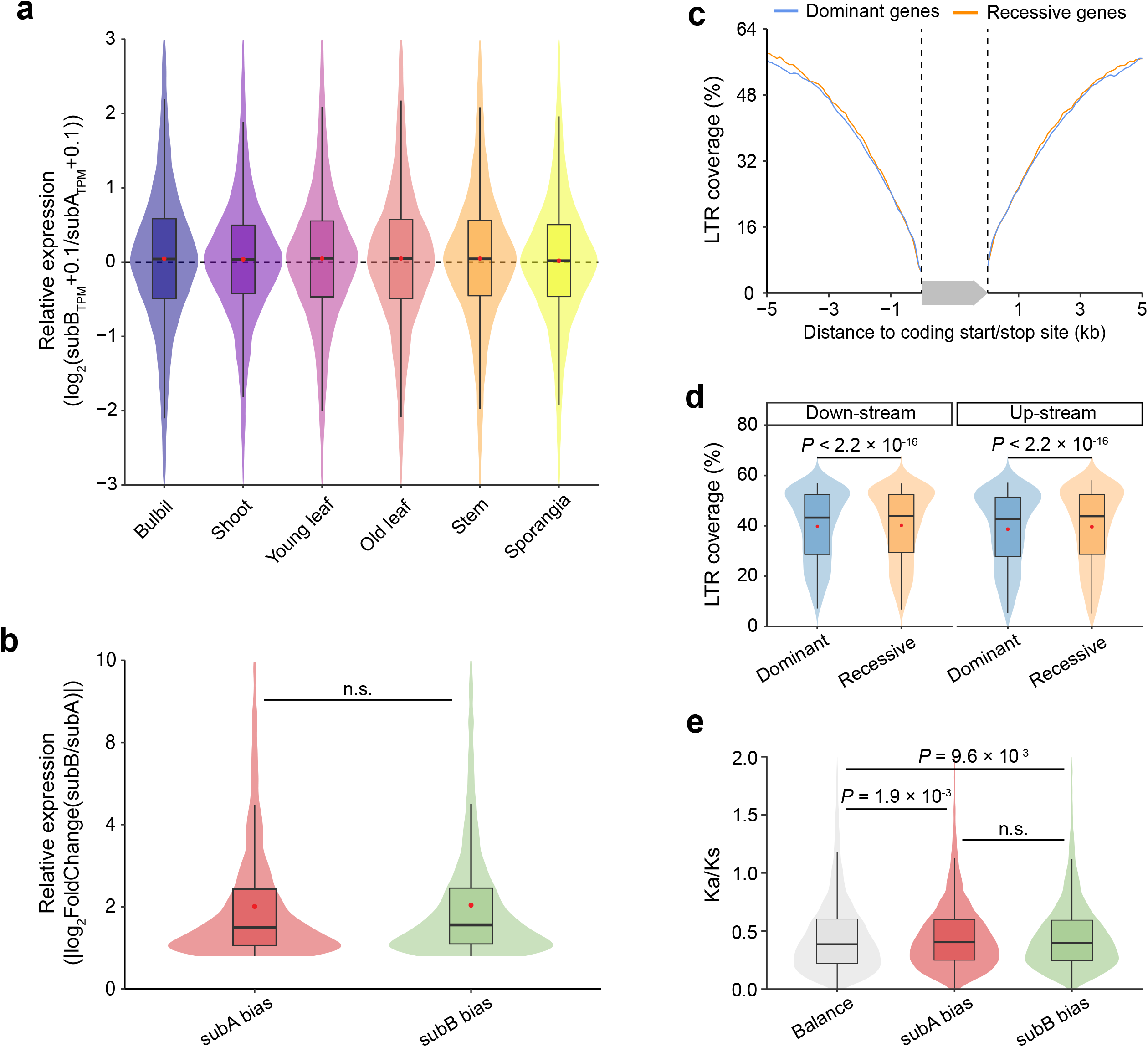
Homoeolog expression bias between two subgenomes of *Huperzia asiatica*. **a,** Relative expression of homoeologous gene pairs in six different tissue types. **b,** Comparisons for absolute value of log_2_FoldChange of homoeologous expression bias (HEB; cutoffs: ‘*P*-value < 0.05 and | log_2_FoldChange | > 1’) gene pairs between two subgenomes of allotetraploid *H. asiatica*. *P*-value estimated by Wilcoxon rank-sum test. ‘n.s’ indicates no significant difference. **c,** Different LTR coverage in the flanking regions between the dominant and the recessive member for each homoeolog gene pair. The two dashed lines and a gray arrow indicate the gene region and orientation, respectively. **d,** Comparisons for LTR coverage the dominant and the recessive member for each homoeologous gene pairs in up- and down-stream of coding region, respectively. *P*-value estimated by Paired Wilcoxon test. **e,** Comparisons for Ka/Ks values of homoeolog gene pairs with expression biased toward subA or subB or with expression balanced between the two subgenomes. *P*-value estimated by Wilcoxon rank-sum test.

Next we investigated possible factors that could give rise to the observed HEB. First we explored the influence of LTR insertions on HEB gene pairs. We compared the LTR coverage in the flanking regions of the dominant homoeolog (with a higher expression) and the recessive homoeolog (with a lower expression), and discovered that the dominant homoeologs had slightly but significantly lower LTR coverage than the recessive counterparts (Fig. 3c,d). This pattern was further corroborated in each of the sampled tissues (Supplementary Fig. 11). We then compared the ratio of nonsynonymous (Ka) to synonymous substitution rates (Ks) in homoeologous gene pairs to test if pairs with HEB experienced different selective pressure. We found that gene pairs with HEB had significantly higher Ka/Ks ratio than those without. On the other hand, when comparing subA- and subB-biased HEB gene pairs, the ratio does not significantly differ (Fig. 3e).

Taken together, our results suggest that neither subgenome has become dominant following allotetraploidization in *H. asiatica*. Relatively few homoeologous gene pairs show HEB and importantly, they do not consistently bias toward subA or subB. The age of the hybrid is obviously important in interpreting our findings. However, although we know that *H. asiatica* subgenomes diverged from the closest known diploid ∼17.8 MYA, the hybridization and/or polyploidization event could take place anywhere between 17.8 MYA and the time we collected our sample. In other words, the lack of a clear subgenome dominance could be due to the recency of the hybrid or speaks to the slow evolutionary nature of *Huperzia* genomes (or the combination of both).

### Highly preserved inter-genomic synteny despite deep divergence

Comparing *H. asiatica* and *D. complanatum* genomes, we were surprised to find a large number of syntenic gene blocks. In total, 11,236 syntenic gene pairs in 1,515 blocks and 11,402 gene pairs in 1,566 blocks were identified between *D. complanatum* genome and *H. asiatica* subA and subB, respectively (Fig. 4a and Supplementary Fig. 12) – these amount to 27-36% of the total proteomes. We can also find multiple *D. complanatum* chromosomes mapping almost continuously to eight *H. asiatica* chromosomes (four from each subgenome; Fig. 4b). Such a high degree of collinearity between the two genomes is unprecedented given that *H. asiatica* and *D. complanatum* diverged around 368 MYA^30^. As a comparison, the extant flowering plants are only about 197.5-246.5 million years old^31^, and yet, synteny is more fragmented between *Amborella* and other angiosperm genomes^32^. Furthermore, the pattern we found here in homosporous lycophytes stands in stark contrast to the lack of synteny between the heterosporous *Isoetes* and *Selaginella* that diverged at around the same time^13^.

**Fig. 4.**
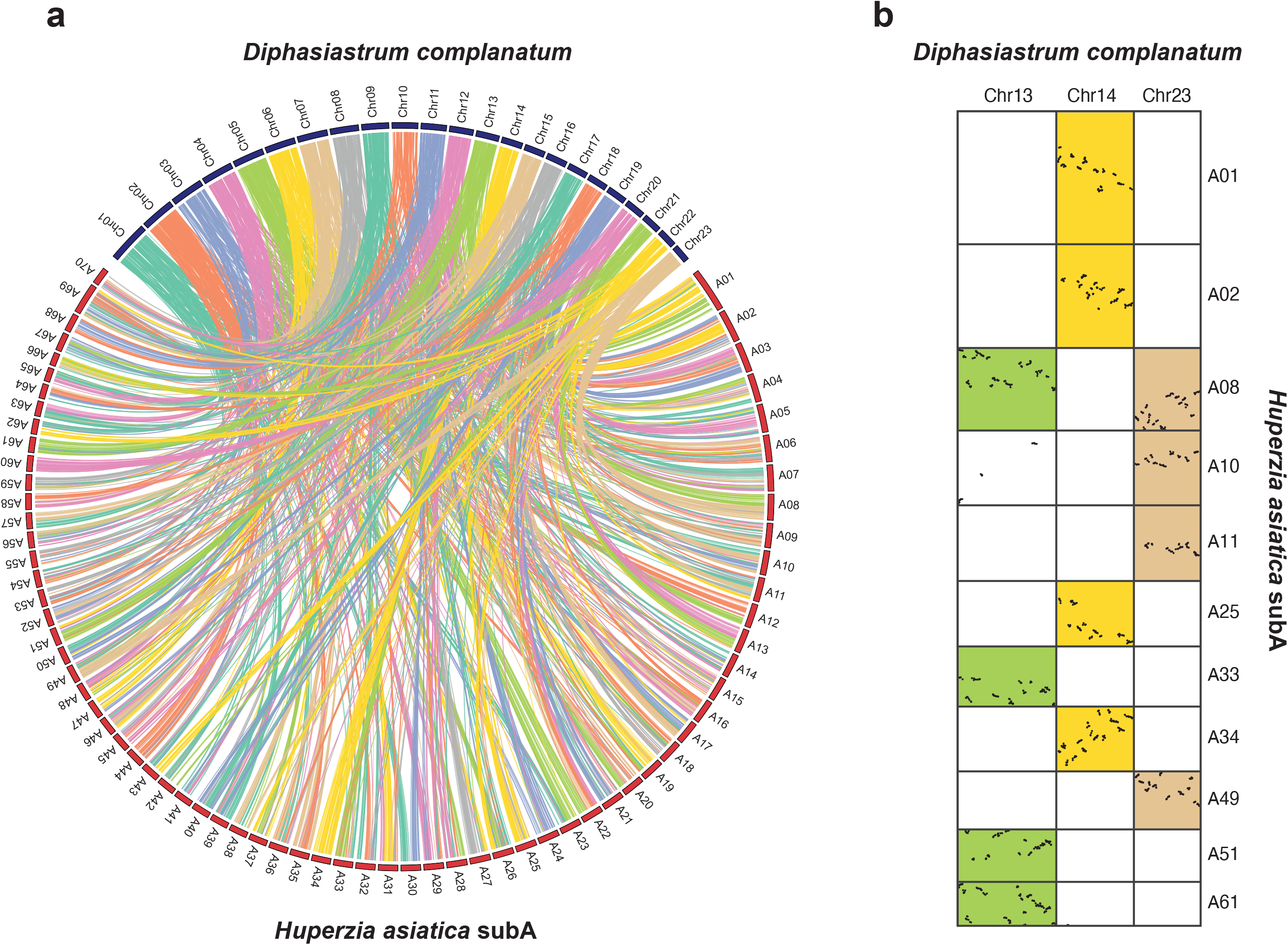
Preserved synteny between distantly related *Huperzia asiatica* and *Diphasiastrum complanatum*. **a,** Syntenic relationship between *H. asiatica* subA and *D. complanatum* genomes. Collinear gene blocks are connected by ribbons. **b,** Syntenic dotplot of representative chromosomes that show a clear 1:4 relationship comparing *D. complanatum* to *H. asiatica* subA.

### Highly preserved intra-genomic synteny despite ancient WGDs

The preservation of synteny can also be observed within each *D. complanatum* and *H. asiatica* genome. In *D. complanatum*, we found 1,422 gene pairs contained in 216 collinear blocks accounting for 9.0% of annotated genes (Fig.1f and Supplementary Fig. 13). In *H. asiatica* subA and subB, we identified 3,284 gene pairs in 338 blocks (17.3% of annotated genes) and 3,778 gene pairs in 429 blocks (18.8% of annotated genes), respectively (Fig.1d,e and Supplementary Fig. 13). The intra-genomic synteny in *D. complanatum* and *H. asiatica* genomes is most likely the result of ancient WGDs. Syntenic depth analysis found a 2:4 relationship between collinear blocks of genes in *D. complanatum* and either of the *H. asiatica* subgenomes (Supplementary Fig. 12), suggesting two WGDs (termed “Huper-α” and Huper-β”) predating the divergence of *H. asiatica* subgenomes and a single WGD (termed “Lyco-α”) in the ancestry of *D. complanatum*.

To further place these WGD events onto phylogeny, we used both synonymous substitutions per site (Ks) and gene tree-species tree reconciliation approaches. Ks analysis of paralogs in *D. complanatum* produced a plot with a single peak centered around Ks = 0.45, suggesting that the Lyco-α event may have occurred following its divergence from *Huperzia* (Fig. 5a and Supplementary Fig. 18). For *H. asiatica*, the Ks plot for each subgenome has a single prominent peak near Ks = 0.2 (Fig. 5b and Supplementary Fig. 19). Although we might expect to see two peaks corresponding to the Huper-α and Huper-β events, given the extraordinarily low orthologous divergence observed between *Huperzia* and *Diphasiastrum* (Fig. 5a,b), it is possible that two WGDs occurring close together would produce overlapping Ks distributions. This hypothesis seems to be supported by a Ks plot restricted to syntenic gene pairs that yields a single peak in each subgenome distributed around Ks = 0.26 (Fig. 5b). Our Ks analyses on related *Huperzia* species revealed an even more complex history of recurrent and independent WGDs throughout the family, a pattern consistent with the high and variable chromosome counts reported. Because of the frequent duplications in this lineage, it is impossible to place WGD events based solely on Ks.

**Fig. 5.**
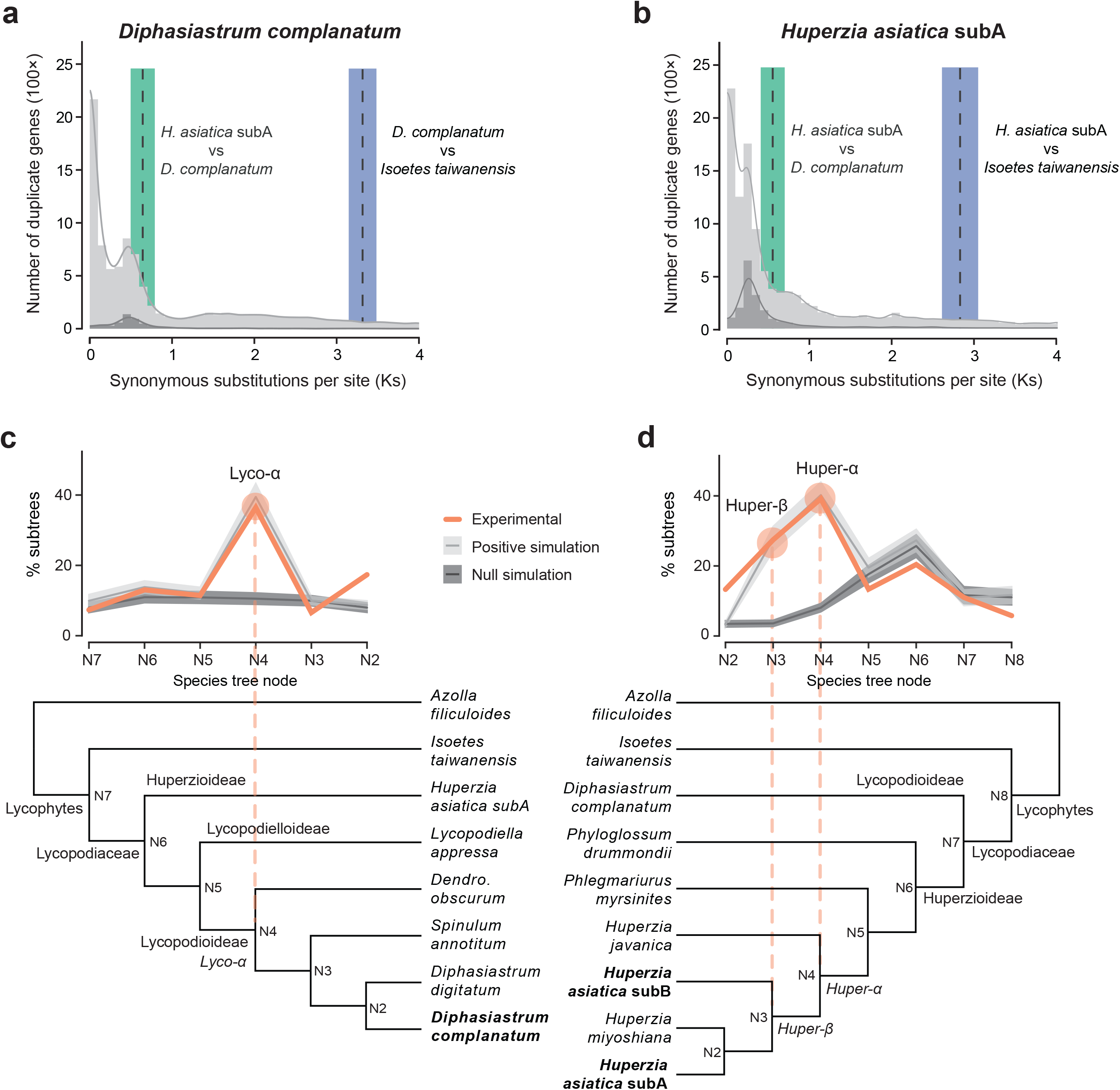
Evidence for ancient WGD in *Huperzia asiatica* and *Diphasiastrum complanatum*. **a,b,** Ks plots of paralogous genes in *D. complanatum* (**a**) and *H. asiatica* subA (**b**). Analysis including all paralogs were shown in light gray, and analysis restricted to collinear gene pairs were shown in dark gray. Dashed vertical lines indicate mean adjusted orthologous divergence for species indicated; shaded areas represent one standard deviation. **c,d,** Summary of MAPs analyses of *D. complanatum* (**c**) and *H. asiatica* subA (**d**). Dark lines in the center of the shaded regions represent average values for null and positive gene tree simulations; shaded areas around each line show the standard deviation for gene tree simulations. Dashed orange lines mark the inferred WGDs.

Using the Multi-tAxon Paleopolyploid Search (MAPS) algorithm, our gene tree-species tree reconciliation approach corroborated that Lyco-α, Huper-α, and Huper-β are all independent events. We found no support for an earlier, shared duplication in the common ancestor of Lycopodiaceae. A MAPS analysis focused on Lycopodioideae (the subfamily including *Diphasiastrum*) placed Lyco-α prior to the divergence of *Dendrolycopodium* and *Diphasiastrum* (Fig. 5c). A separate analysis focused on the Huperzioideae implicated Huper-α and Huper-β occurred following the divergence of *Huperzia* from its sister genus *Phlegmariurus* (Fig. 5d and Supplementary Fig. 20).

Based on their phylogenetic placements, the three WGD events we uncovered here are all ancient. Lyco-α in particular is likely at least 139 MYA, given it predates the divergence between *Diphasiastrum* and *Dendrolycopodium*^30^. Despite the antiquity of this duplication, there are still 9% of the annotated genes retained in synteny. Such a slow rate of genome rearrangement echoes what we reported above regarding the divergence between *D. complanatum* and *H. asiatica*.

### Reciprocal fractionation may obscure intragenomic synteny in homosporous lineages

Despite a predominant 2:4 relationship between syntenic blocks of genes in *D. complanatum* and *H. asiatica* subgenome (Supplemental Fig. 12), we found relatively few genes showing the expected 2:2 relationship within the *H. asiatica* subgenome (Supplemental Fig. 13). While this may at first seem counterintuitive, it can be explained by the process of differential fractionation of genes in each species following WGD. Immediately following duplication, species begin to undergo the process of diploidization, by which diploid patterns of gene expression and inheritance are restored through gene fractionation and chromosomal rearrangement. In heterosporous species, these processes are tightly coupled with many genes being lost rapidly via illegitimate recombination^33^. However, high numbers of chromosomes in homosporous lineages of ferns and lycophytes suggest that such rapid, large scale losses and rearrangements occur less frequently^34,35^. Instead, the primary processes driving diploidization in homosporous plants likely involve silencing and eventual loss of individual genes^34,36^. Thus, through the reciprocal loss of paralogs along homoeologous chromosomes, it is possible to produce two sets of homoeologous chromosomes with a relatively high degree of intergenomic synteny despite exhibiting little to no intragenomic synteny (Supplemental Fig. 14). If this initial gene loss was followed by a second WGD in *H. asiatica*, we would then expect a 1:1 relationship to be restored between newly duplicated chromosomes assuming that there has been less fractionation following this more recent WGD. In fact, when we zoom in on scaffolds that exhibit a 2:4 relationship between *Huperzia* and *Diphasiastrum*, this is precisely what we see (Supplemental Figs. 15-17). Large overlapping blocks are anchored by distinct genes scattered across the same region of the scaffold with relatively low levels of intragenomic synteny relative to intergenomic synteny. Despite this process of diploidization by fractionation, the retention of long range synteny both within and between these species is remarkable given the timescales involved and unlike anything previously described in heterosporous plants. Furthermore, this pattern seems to corroborate the hypothesis that diploidization in homosporous lineages is largely driven by silencing and fractionation of individual genes rather than large-scale structural changes^34,35^.

### Highly preserved synteny is linked to decelerated rates of nucleotide substitution

The collinearity found in *H. asiatica* and *D. complanatum*, both within and between genomes, are in stark contrast to sister, heterosporous lineages (*Isoetes*^13^ and *Selaginella*^12^). Our placement of the Lyco-α WGD event prior to the divergence of *Diphasiastrum* and *Dendrolycopodium* renders intraspecific synteny within *D. complanatum* on par with those recently found in tree fern *Alsophila spinulosa*^20^. Similarly, while limited examples of ancient inter- and intraspecific collinearity have been reported in species of fish^37,38^ and moss^39,40^, divergence time estimates for *Diphasiastrum* and *Huperzia* are considerably older^30^. Surprisingly, given their great evolutionary distance, orthologous divergence between *H. asiatica* and *D. complanatum* is relatively low (Ks ∼ 0.5-0.7; Fig. 5a,b) implying a slow rate of substitution. As previously described for tree ferns^20^, we found decelerated rates of substitution in Lycopodiaceae compared to heterosporous lycophytes *Selaginella* and *Isoetes*. Most of the gene families examined show signs of deceleration and 22-31% of the comparisons are significant (Fig. 6). These results, combined with those from the tree fern *A. spinulosa* indicate that the gradual processes of chromosomal rearrangement and fractionation may be correlated with relatively slow rates of substitution in homosporous seed-free lineages.

**Fig. 6.**
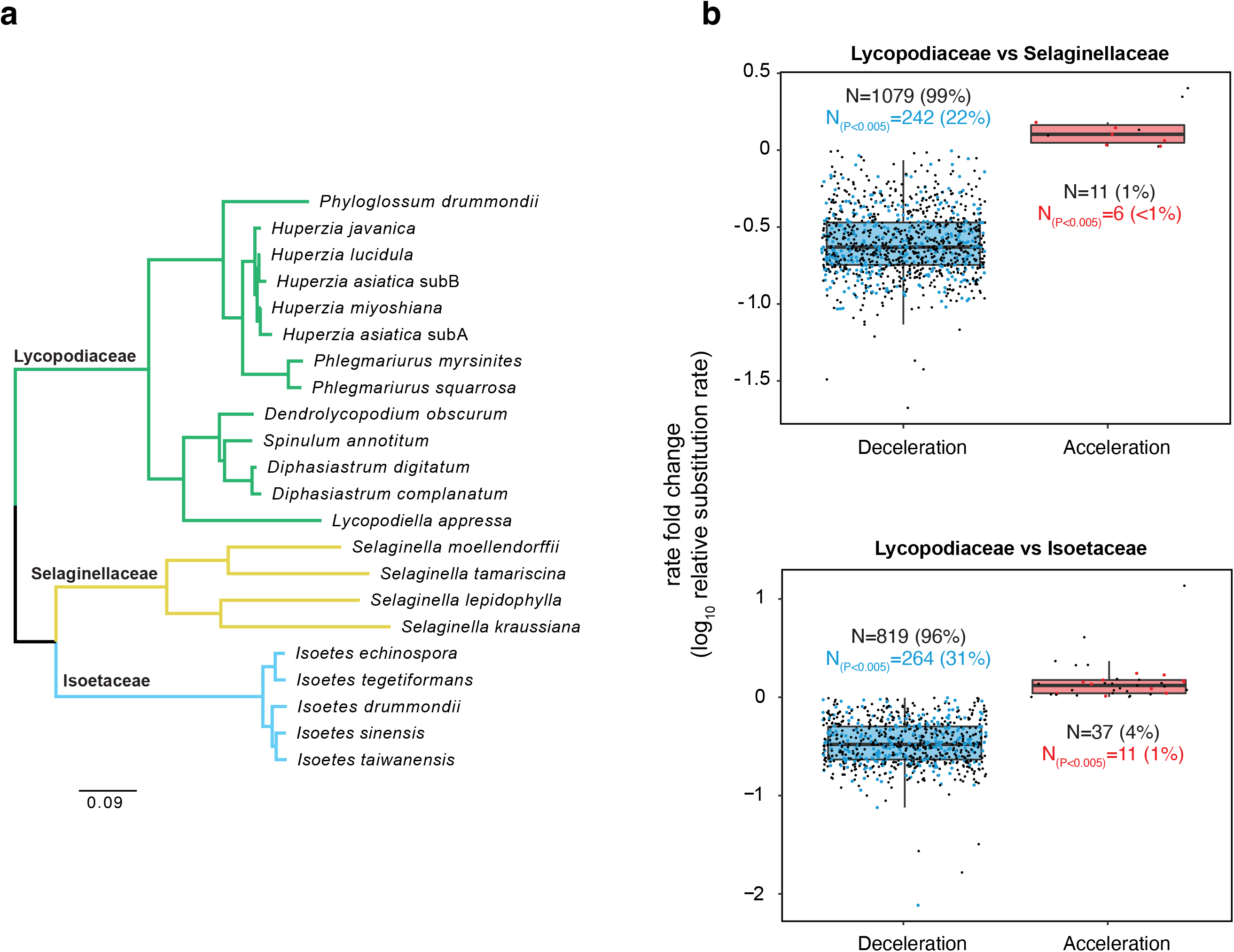
Lycopodiaceae exhibits lower substitution rates compared to heterosporous lycophyte lineages. **a,** Phylogenetic tree with branch colors denoting lycophyte families demonstrating evolutionary relationships within species used to analyze substitution rates. **b,** Boxplots showing differences in substitution rates between Lycopodiaceae and species from heterosporous lineages.

In summary, our study suggests that homosporous lycophytes likely have a contrasting mode of genome evolution compared to heterosporous lycophytes and sheds new light on the process of diploidization following WGD. Homosporous lineages are known to harbor exceptionally large genomes with high chromosome counts relative to other vascular plants. This contrast has puzzled botanists for over half the century^4,35,41,42^ and no mechanistic explanation has been widely accepted so far. Our research reveals a number of factors contributing to the markedly larger genome sizes of homosporous lycophytes, including an abundance of TE insertions, extreme intron length, and multiple rounds of WGD. Importantly, we found that the rates of molecular evolution decelerated, specifically those of substitution, fractionation, and chromosomal rearrangement. It is tempting to generalize our findings to all homosporous vascular plants, especially considering that similar findings were reported from the homosporous fern *Alsophila spinulosa*^20^. However, the more recent publications of *Ceratopteris richardii*^16^ and *Adiantum capillus-veneris*^17^ genomes suggest the same might not be true in other homosporous ferns, where genome fractionation appears to be rapid. While the current sampling of homosporous genomes is too narrow to give us a complete picture, our study has nevertheless identified a possible genomic contrast between homosporous and heterosporous species. This testable hypothesis will set the stage for future comparative studies on seed-free genomes.

## Methods

### Plant materials

The *Huperzia asiatica* plants for de novo assembly were collected from Changbai Mountain National Nature Reserve in Antu County, Jilin Province, China. The *Diphasiastrum complanatum* plants for de novo assembly were collected from Yilan County, Taiwan.

### Genome size estimation

K-mer and flow cytometry methods were used for genome size estimation. Jellyfish^43^ was performed to generate a 21-mer frequency distribution, which was then analyzed in GenomeScope^44^ to estimate the genome size. For flow cytometry analysis, a modified protocol from published procedure^45^ was utilized to assess the genome size. Briefly, chopped fresh shoots were incubated in 1 mL of LB01 solution (Coolaber Science & Technology Co., Ltd, Beijing, China) for 30 min to release nuclei. Nuclei suspension was filtered through a 40 μm of cell strainer, followed by the addition of 20 μg mL^-1^ Rnase A (Coolaber Science & Technology Co., Ltd, China) and 20 μg mL^-1^ propidium iodide (PI) (Coolaber Science & Technology Co., Ltd, China) and incubated on ice for 30 min in the dark. The fluorescence intensity of PI-stained nuclei was determined using the flow cytometer CytoFLEX (Beckman Coulter, USA) with a flow rate of 10 μL min^-1^ and three biological replicates. For *H. asiatica* and *D. complanatum*, we used *Taxus chinensis* (1C = 10.23 Gb) ^46^ and *Zea mays* Mo17 (1C = 2.15 Gb) ^47^ as the standards, respectively. Fluorescence histograms were analyzed with CytExpert (a special analysis software for CytoFLEX) to infer the nuclear DNA.

### Genome sequencing

We sequenced *Huperzia asiatica* and *Diphasiastrum complanatum* using a combination of sequencing technologies, including long-read sequencing from Pacific Biosciences (PacBio), Illumina short-read sequencing, and chromosome conformation capture using Hi-C sequencing.

The PacBio SMRTbell libraries of *H. asiatica* and *D. complanatum* were constructed with high molecular weight (HMW) genomic DNA using the PacBio SMRTbell express template prep kit 2.02. The libraries were size-selected on a BluePippin system for molecules longer than 15 Kb, followed by primer annealing and the binding of SMRTbell templates to polymerases by using the DNA/Polymerase Binding Kit. The long reads sequencing was performed on the PacBio Sequel II platform with ‘Continuous Long Read (CLR) Sequencing’ mode. We generated ∼75× PacBio CLR reads (589.68 Gb with average read size 18.02 Kb) for *H. asiatica* and ∼139× (242.47 Gb with average read size 16.15 Kb) for *D. complanatum* (Supplementary Table 9).

For Illumina short-read sequencing, high-quality genomic DNA of each sample was extracted by using a DNeasy Plant Mini Kit (Qiagen, United States). The DNA sequencing libraries were constructed following the procedures provided by the Nextera XT DNA Library Prep Kit (Illumina, Inc., San Diego, CA, USA). The insert size of each library was verified by the Agilent Bioanalyzer 2100 system. Library preparations of DNA were sequenced on the Illumina NovaSeq 6000 platform (Illumina Inc, USA). For *D. complanatum*, 104.21 Gb for a total of ∼65× of Illumina reads were produced, and 416.22 Gb Illumina reads were yielded for *H. asiatica* with a total coverage of ∼53× (Supplementary Table 9).

Hi-C libraries were constructed from fresh shoots of *H. asiatica* and *D. complanatum* according to previously published procedure^48,49^. Briefly, the plant tissues were fixed with formaldehyde, lysed, and cross-linked DNA digested with HindIII over-night. The sticky ends of the digested fragments were biotinylated, diluted, and ligated randomly. Chimeric fragments representing the original cross-linked long-distance physical interactions were processed into paired-end sequencing libraries. The sequencing of these libraries was carried out with Illumina NovaSeq 6000 platform with 2×150 bp reads.

### RNA-Seq

Total RNA was extracted from *Huperzia asiatica*, *H. miyoshiana*, *H. serrata*, *H. lucidula*, and *Diphasiastrum complanatum* by using the RNeasy Plant Mini kit (Qiagen). The integrity of the RNA was determined with an Agilent 4200 Bioanalyzer system (Agilent Technologies, United States). High quality RNA from each sample was used for cDNA library construction using ReverTra Ace (TOYOBO, Osaka, Japan) with oligo (dT) primers following the user manual. The PCR products obtained were purified (AMPure XP system) and library quality was assessed on an Agilent Bioanalyzer 2100 system (Agilent Technologies, United States). Library preparations of RNA were sequenced on the Illumina NovaSeq 6000 platform (Illumina Inc, USA) or MGISEQ-2000 platform (BGI-Shenzhen, China) with 150 bp paired-end read length (Supplementary Table 10). Three biological replicates for shoot, bulbil, young leaf, old leaf, stem, and sporangia of *H. asiatica* were sampled.

### Iso-Seq

Full-length cDNA was synthesized with 1.0 μg RNA from *Diphasiastrum complanatum* using the SMARTer PCR cDNA Synthesis kit (Clontech). The first-strand cDNA was amplified with PrimeSTAR GL DNA polymerase (Clontech) to generate double-stranded cDNA. Double-stranded cDNA was purified with non-size selection AMPure PB beads (PacBio). The amplified cDNA was ligated with blunt-end PacBio sequencing adaptors using SMRTbell Template Prep Kit. The ligated products were treated by exonuclease to remove un-ligated products and purified by AMPure PB beads (PacBio). The sequencing was performed on the PacBio Sequel II platform.

### Genome assembly and subgenome phasing

The 589.68 Gb PacBio CLR subreads of *Huperzia asiatica* were first self-corrected using CANU^50^ with parameters ‘minReadLength=1000 minOverlapLength=500 corOutCoverage=120 corMinCoverage=2’ and about 480.30 Gb (∼61×) corrected subreads were obtained. After trimming the corrected subreads, the assembly was performed using ‘canu -assemble’ with parameter ‘correctedErrorRate=0.045’. Then, we performed Purge_haplotigs program^51^ to filter the redundant contigs from the primary assembly. The resulting contigs were polished with one round of alignment with PacBio CLR subreads by using Racon^52^ and two rounds of alignment with Illumina short reads by using NextPolish^53^, respectively.

The Hi-C Illumina reads from *H. asiatica*, were aligned to the contig sets of *H. asiatica*, respectively, with Juicer^54^, and the scaffolding was performed with 3D-DNA^55^. The Juicebox Assembly Tools^56^ were then used for the manual correction of the connections. For contamination removal, the Hi-C data could theoretically distinguish DNA sequences originating from different organisms because cross-linking occurred within the nuclei. We reasoned that parts of the unanchored sequences could come from non-streptophyte organisms. Hence, the unanchored scaffolds or contigs were used to conduct a MEGABLAST search against NCBI NT (https://www.ncbi.nlm.nih.gov/) database to remove the non-streptophyte originated sequences or organellar DNA. A total of approximately 65.86 Mb of sequences were removed from the final assembly of *H. asiatica*.

Through Hi-C interaction heatmap, we could determine the pairwise relationship between the homoeologous chromosomes of allotetraploid *H. asiatica*. Next, we partitioned each of homoeolog pairs into two sets of subgenomes using SubPhaser^57^ based on the subgenome-specific k-mers. First, 15-bp sequences (15-mer) counting was carried out via Jellyfish^43^ in all the chromosomes. Second, for each 15-mer whose frequency was larger than 200 (‘-min_freq 200’) was selected for further analysis. The 15-mers, whose abundance were at least twice (‘-min_fold 2’) in one member than that in another in the homoeolog chromosome pairs, were identified as subgenome-specific. Third, the k-means algorithm was performed for clustering all the chromosomes into two subgenomes based on the different abundance of subgenome-specific 15-mers. Forth, principal component analysis (PCA) and 15-mer abundance heatmap were used to verify the phasing results. Then, these subgenome-specific 15-mers were mapped against the genome via a substring match procedure, and the 15-mer enrichment degree in a 1 Mb window was determined by Fisher’s exact test (*P*-value < 0.05). Minimap2^58^ is used to search homoeologous blocks between homoeologous chromosome pairs.

For *Diphasiastrum complanatum*, the main assembly consisted of 139× of PacBio coverage (16,154 bp average read size), and was assembled using MECAT^59^, with the resulting sequence being polished using ARROW^60^. 8 misjoins in the assembly were identified using Hi-C data. Contigs terminating in significant telomeric sequences were properly oriented in the assembly. A total of 3,482 joins were applied to the assembly to form the final assembly consisting of 23 chromosomes with 83.08% of the assembled sequence in chromosomes. Adjacent alternative haplotypes were identified on the joined contig set. Althap regions were collapsed using the longest common substring between the two haplotypes. A total of 200 adjacent alternative haplotypes were collapsed in the assembly. Chromosomes were numbered from smallest to largest with the p-arm oriented to the 5’ end, and the resulting sequence was screened for retained vector and/or contaminants. Heterozygous snp/indel phasing errors were corrected using the 139× PacBio data. A total of 30,496 heterozygous SNPs/INDELs were corrected. Additionally, 43,307 homozygous SNPs and INDELs were corrected in the release sequence using ∼62× of Illumina reads.

### Genome annotation

For transposable elements (TE) annotation of *Huperzia asiatica*, the TE library was obtained using Extensive de novo TE Annotator (EDTA v1.9.4) pipeline^61^, which integrated multiple predictors, including LTR_FINDER_parallel^62^, parallel modified LTRharvest^63^, LTR_retriever^64^, TIR-Learner^65^, MITE-Hunter^66^, and HelitronScanner^67^. To enable better parallel computation and accelerate the annotation for genomes larger than 3 Gb, we split the genome assembly into four equally sized parts according to developers’ suggestion (https://github.com/oushujun/EDTA/issues/61). Briefly, the TE library of each part from the genome was independently predicted by running the ‘EDTA.pl’ script in the EDTA package with parameters ‘--sensitive 1 --anno 1’. The four TE libraries were integrated by using the ‘make_panTElib.pl’ script in the EDTA package to generate a high-quality TE library of *H. asiatica*. The final TE library was then used to annotate each part of the genome by RepeatMasker (http://www.repeatmasker.org) with parameters ‘-q -no_is -norna -nolow -div 40 -cutoff 225’. To annotate the intact TEs in of *H. asiatica*, the homology-based TE annotation of each part from the genome (the outputs from RepeatMasker) was fed to ‘EDTA.pl’ script with parameters ‘--evaluate 1 --anno 1 --step final’, respectively. The repeat-masked genome was generated by running ‘make_masked.pl’ script in the EDTA package with parameters ‘-maxdiv 40 -minscore 300 -minlen 80’. To identify tandem repeats in the genomes, TRF^68^ was used with parameters ‘1 1 2 80 5 200 2000 -d -h’. Annotation of protein-coding genes of *H. asiatica* was based on *ab initio* gene predictions, transcript evidence and homoeolog protein evidence, all of which were performed in the MAKER^69^. In brief, transcript evidence was supplied in the form of a de novo assembled transcriptome produced with TRINITY^70^ and a reference-based assembly generated by StringTie^71^. The proteome data of *Anthoceros agrestis* Bonn^19^, *Marchantia polymorpha*^72^, *Physcomitrium patens*^18^, *Azolla filiculoides*^73^, *Salvinia cucullata*^73^, *Selaginella moellendorffii*^10^, *S. lepidophylla*^12^, *Isoetes taiwanensis*^13^, *Amborella trichopoda*^32^, *Oryza sativa*^74^, *Arabidopsis thaliana*^75^, and Swiss-Prot (https://www.uniprot.org/downloads/) database as the protein evidence. The first round of MAKER was implemented with the transcript and protein evidence. To train AUGUSTUS^76^ and GENEMARK^77^, BRAKER2^78^ was applied with the transcript data from aligned RNA-Seq bam files produced by HISAT2^79^. SNAP^80^ was trained under MAKER with two iterations. For the second round of MAKER, the results from AUGUSTUS, GENEMARK, SNAP were fed into MAKER again along with all other data to produce synthesized gene models.

For genome annotation of *Diphasiastrum complanatum*, transcript assemblies were made from ∼1 billion pairs of 2×150 stranded paired-end Illumina RNA-seq reads using PERTRAN (https://www.osti.gov/biblio/1241180). About 21 million PacBio Iso-Seq CCSs were corrected and collapsed by genome guided correction pipeline (Shu, unpublished) to obtain ∼859,000 putative full-length transcripts. A total of 722,481 transcript assemblies were constructed using PASA^81^ from RNA-Seq transcript assemblies above. Loci were determined by transcript assembly alignments and/or EXONERATE^82^ alignments of proteins from *Marchantia polymorpha*^72^, *Physcomitrium patens*^18^, *Selaginella moellendorffii*^10^, *Amborella trichopoda*^32^, *Oryza sativa*^74^, *Arabidopsis thaliana*^75^, *Glycine max*^83^, *Sorghum bicolor*^84^, *Setaria viridis*^85^, *Aquilegia coerulea*^86^, *Solanum lycopersicum*^87^, *Vitis vinifera*^88^, *Nymphaea colorata*^89^, *Papaver somniferum*^90^, and Swiss-Prot (https://www.uniprot.org/downloads/) proteomes to repeat-soft-masked *D. complanatum* genome using RepeatMasker (http://www.repeatmasker.org) with up to 2kp extension on both ends unless extending into another locus on the same strand. Repeat library consists of de novo repeats by RepeatModeler (https://github.com/Dfam-consortium/RepeatModeler) on *D. complanatum* genome and repeats in RepBase. Gene models were predicted by homoeology-based predictors, FGENESH+^91^, FGENESH_EST (similar to FGENESH+, but using EST to compute splice site and intron input instead of protein/translated ORF), and EXONERATE, and PASA assembly ORFs (in-house homology constrained ORF finder). The best scored predictions for each locus are selected using multiple positive factors including EST and protein support, and one negative factor: overlap with repeats. The selected gene predictions were improved by PASA. Improvement includes adding UTRs, splicing correction, and adding alternative transcripts. PASA-improved gene model proteins were subject to protein homology analysis to above mentioned proteomes to obtain Cscore and protein coverage. Cscore is a protein BLASTP score ratio to MBH (mutual best hit) BLASTP score and protein coverage is highest percentage of protein aligned to the best of homologs. PASA-improved transcripts were selected based on Cscore, protein coverage, EST coverage, and its CDS overlapping with repeats. The transcripts were selected if its Cscore is larger than or equal to 0.5 and protein coverage larger than or equal to 0.5, or it has EST coverage, but its CDS overlapping with repeats is less than 20%. For gene models whose CDS overlaps with repeats for more than 20%, their Cscore must be at least 0.9 and homology coverage at least 70% to be selected. The selected gene models were subject to Pfam analysis and gene models whose protein is more than 30% in Pfam TE domains were removed and weak gene models. Incomplete gene models, low homology supported without fully transcriptome supported gene models and short single exon (< 300 BP CDS) without protein domain nor good expression gene models were manually filtered out.

The genome features of *Huperzia asiatica* and *Diphasiastrum complanatum* were visualized by Circos with the non-overlapped sliding windows set as 1 Mb and 500 Kb, respectively. The average gene length for *H. asiatica* subA is 20.78 Kb and the maximum gene length is 57.25 Kb. With a sliding window size of 1 Mb, we have on average ∼15 genes per sliding window (average length of intergenic region is 45 Kb) in *H. asiatica* subA. For *D. complanatum*, the average gene length is 12.90 Kb and the maximum gene length is 270.54 Kb. With a sliding window size set to 500 Kb, on average ∼12 genes are covered per sliding window (average length of intergenic region is 30 Kb).

The annotation qualities of *Huperzia asiatica* and *Diphasiastrum complanatum* were tested by BUSCO^92^ with viridiplantae_odb10 database. Functional annotations of the protein-coding sequences were carried out by using BLASTP searches against entries in both NCBI NR (https://www.ncbi.nlm.nih.gov/) and Swiss-Prot (https://www.uniprot.org/) databases with cutoff ‘-e-value 1.0×10^-10^’. The prediction of conserved domains for the coding genes was performed via InterProScan^93^. The annotations of the GO terms (http://geneontology.org/) and KEGG pathways (https://www.genome.jp/kegg/) of the coding genes were obtained from eggNOG-mapper^94^.

### Origin of the allotetraploid *Huperzia asiatica*

To infer the age of the subgenomes and allotetraploid *Huperzia*, an orthologous set of protein sequences from 12 Lycopodiaceae genome or transcriptome assemblies, including *H. asiatica* subA, *H. asiatica* subB, *H. selago*^9^, *H. miyoshiana*, *H. lucidula*^9^, *H. serrata*, *H. javanica*^95^, *Phlegmariurus myrsinites*^9^, *P. squarrosa*^9^, *Phylloglossum drummondii*^9^, *Lycopodiella appressa*^9^, and *Diphasiastrum complanatum*, was generated based on OrthoFinder^96^ with *D. complanatum* as outgroup. We identified 1,233 low-copy orthologous groups (at most two gene copies of each orthologous group for each species). The longest isoform for each protein was retained. We then reconstructed the phylogenetic tree by concatenation-based and coalescence-based methods as described above. The phylogenetic tree was used as the input for mcmctree^97^ to compute a time tree with a fossil constraint of Tracheophyta set to 421-451 MYA^31^. The program discarded the first 5,00,000 iterations (parameter ‘burnin’), and then it sampled every 50 iterations (parameter ‘sampfreq’) until 1,000,000 samples were collected (parameter ‘nsample’). The resulting trace file of the MCMC program was evaluated using Tracer^98^ to ensure convergence; the effective sample size (ESS) of all parameters were confirmed to be at least 200.

To determine the maternal progenitor of *Huperzia asiatica*, we constructed the chloroplast phylogeny. First, we assembled the chloroplast genomes of *H. asiatica*, *H. miyoshiana*, *H. serrata*, *H. lucidula*, *H. selago*, and *Diphasiastrum complanatum*. Briefly, we downloaded the short-read data of *H. selago* from NCBI (https://sra-pub-run-odp.s3.amazonaws.com/sra/ERR5554789/ERR5554789), and generated the sequencing data for the remaining five species based on either Illumina platform (NovaSeq 6000) or BGI platform (MGISEQ-2000) (Supplementary Table 9). All the short-read data from the six species were filtered by using fastp^99^, aligned to the chloroplast database (containing all published chloroplast genomes from NCBI by the date November 26, 2019) with BWA^100^, and then selected chloroplast reads with Picard (https://broadinstitute.github.io/picard/). The selected chloroplast reads were assembled by Spades^101^ with default parameters and the output scaffolds (GFA file) were imported into Bandage^102^ to generate the final chloroplast genome for each species. The six chloroplast genome sequences were aligned with MAFFT^103^ and trimmed using trimAl^104^ to remove the poorly aligned positions. Finally, the chloroplast phylogeny was reconstructed based on the aligned sequences using RAxML^105^ with 1,000 bootstrap replicates.

### Synteny analysis

Syntenic blocks of genes were identified within and between the *Diphasiastrum complanatum* genome and *Huperzia asiatica* subgenomes using MCSCAN^106^. Using CDS and annotation gff3 files as input data, we used MCSCAN’s ‘jcvi.compara.catalog ortholog’ function to identify and visualize intra-/inter-genomic syntenic regions. The default C-score of 0.7 is used to filter low-quality hits. The syntenic blocks were filtered using ‘jcvi.compara.synteny screen’ function with parameter of ‘-minspan = 30’. Synteny pattern was detected using ‘jcvi.compara.synteny depth’ function.

### Heterozygosity estimation

To calculate the genome-wide heterozygosity (GWH) of *Huperzia* species and *Diphasiastrum complanatum*, we first generate draft genome assemblies (consensus sequences) of *H. miyoshiana*, *H. serrata*, and *H. lucidula*. The raw Illumina short reads were filtered by using fastp^99^ with default parameters, mapped onto their corresponding subgenome references of *H. asiatica* with BWA^100^. We sorted the mapping bam files using SAMtools^107^ and then generated their genomes (consensus sequences) using the doFasta module of ANGSD^108^ with parameters ‘-doFasta 2 - doCounts 1 -minMapQ 30 -minQ 20’. A minimum depth (‘-setMinDepth’) of 8, 10 and 30 was set for *H. miyoshiana* (∼55×, relative to *H. asiatica* subA), *H. serrata* (∼64×, relative to *H. asiatica* subB) and *H. lucidula* (∼144×, relative to *H. asiatica* subB), respectively, based on their different sequencing depth. Then the GWH of each species was estimated using ANGSD based on a Site Frequency Spectrum (SFS). The saf.idx file of each species was generated by the doSaf function in ANGSD with parameters of: a minimum mapping quality of 30 (‘-minMapQ 30’), a minimum base quality of 20 (‘-minQ 20’), and an estimation of genotype likelihoods by SAMtools (‘-GL 1’), respectively, for the de novo assembled genomes of *H. asiatica* and *D. complanatum* as well as the consensus genomes of *H. miyoshiana*, *H. lucidula* and *H. serrata*. The SFS for each chromosome from species was estimated using the realSFS packaged in ANGSD and the ratio of heterozygous sites/total sites was calculated as the heterozygosity for each chromosome.

### Demographic inference

The Pairwise Sequentially Markovian Coalescent (PSMC) method^23^ was used to infer the population size history of *Huperzia asiatica* and *Diphasiastrum complanatum*. For each species, 30× genome coverage of the illumina paired-end reads was extracted and aligned to the genome (consensus sequences) using BWA. PCR duplicates were removed using SAMtools^107^. Heterozygous biallelic SNPs were called using SAMtools and consensus sequences were generated using BCFtools^109^. The consensus sequence was fed to the PSMC. All of the parameters used for the PSMC program were at default with the exception: the mutation rate of 6.0×10^-10^ and the generation time (-g) of 15^110,111^. The consistency of the PSMC results was tested by performing 100 bootstrap replicates. We calculated the Ks values of homoeologous gene pairs between *H. asiatica* subgenomes and the peak of the Ks is 0.04. According to the formula T = Ks/2r and the divergence time between *H. asiatica* subA and subB (∼33.9 MYA), we estimated a mutation rate (r) of 6.0×10^-10^ substitutions per synonymous site per year for further analysis.

### Subgenome homoeolog expression

Raw transcriptome reads of six tissue types, including shoot, bulbil, young leaf, old leaf, stem, and sporangia, from allotetraploid *Huperzia asiatica* were filtered using fastp^99^ to remove adapters, mapped to the genome reference using HISAT2^79^ with default settings, and then sorted by SAMtools^107^. Transcripts were quantified in transcripts per million (TPM) from RNA-seq alignments using StringTie^71^. The built-in script ‘preDE.py’ in StringTie was used to convert the quantification results into a count matrix. Based on the results of syntenic analysis, we identified 18,264 homoeologous gene pairs between two subgenomes of allotetraploid *H. asiatica*. Homoeologous expression bias (HEB) gene sets were identified between all the homoeologous gene pairs of two subgenomes using the DESeq2^112^ package for all the six tissues. The Wald test built in the DESeq2 was used to test whether the homoeologous gene pairs have significantly different (i.e. biased) expression levels. Here, we set two cutoffs as follows: (i) the absolute value of ‘log2FoldChange’ between the homoeologous gene pairs was higher than 1; (ii) the false discovery rate (the corrected *P*-value) was less than 0.05.

### LTR coverage in the flanking regions of homoeologous expression bias (HEB) gene pairs

For homoeologous expression bias (HEB) gene pairs in allotetraploid *Huperzia asiatica*, two 5 kb sequences in their 5’ and 3’ regions were identified as their flanking sequences. A 100 bp sliding window made by BEDTools^113^ ‘makewindows’ sub-command with a 10 bp step was used to screen in the 5’ and 3’ flanking regions. In each window, the coverage of LTR sequences was calculated for the 5’ and 3’ flanking regions of each gene by using sub-command ‘coverage’ built in BEDTools, and then the average of coverage was computed for dominant and recessive members for each HEB gene set.

### Ancient whole genome duplication (WGD)

Paralogous Ks estimation and fitting of generalized mixture models were conducted with the WGD package^114^ using the ksd and mix commands. In addition to estimating paralogous and orthologous Ks within and between *Diphasiastrum complanatum* and *Huperzia asiatica* subgenomes, Ks analysis was conducted for 10 other species in Lycopodiaceae, whose transcriptomic data was available. Due to the relatively low Ks divergence between *Huperzia* and *Diphasiastrum* as well as prior evidence of a high degree of rate heterogeneity among lycophytes in general^30^, we attempted to clarify the relative timing of WGD events by adjusting orthologous divergence using KsRates^115^. KsRates adjusts orthologous Ks peaks using an outgroup to assign branch-specific contributions to Ks and rescaling those values to the Ks timescale of a particular focal species. Thus, the adjustment of proximal Ks values relies on initial comparison to the outgroup. In this case, due to the phylogenetic placement of *Huperzia* relative to *Diphasiastrum* we are forced to use heterosporous species of lycophytes as our closest outgroups. Unfortunately, highly divergent outgroups often exhibit relatively flat Ks distributions with high values approaching saturation of synonymous substitutions. This results in large error rates for initial estimates of branch specific Ks that are then carried forward into rate adjustment. This issue of deep divergence is further exacerbated in lycophytes by high levels of substitution rate heterogeneity between these groups. As a result, Ks and even rate adjusted Ks may not be a particularly informative metric for determining the relative timing of WGD events in this group. That being said, rate adjustment did at least appear to confirm that duplications in both groups occurred following their divergence. We thus turned to phylogenetic reconciliation in MAPs to try and more accurately place the hypothesized WGDs in each lineage.

In order to conduct phylogenetic reconciliation, three separate datasets were compiled using the protein coding sequences from *Azolla filiculoides*^9^, *Dendrolycopodium obscurum*^9^*, Diphasiastrum complanatum*, *D. digitatum*^9^, *Huperzia asiatica*, *H. javanica*^95^, *H. lucidula*^9^, *H. miyoshiana*, *Isoetes taiwanensis*^13^*, Lycopodiella appressa*^9^*, Phlegmariurus myrsinites*^9^, and *Phyloglossum drummondii*^9^. OrthoFinder^96^ was used for clustering, alignment, and construction of gene trees with the ‘-m MSA’ flag. Gene trees from OrthoFinder were filtered with a custom python script to retrieve trees with at least one gene from each taxon in the analysis and modified to remove branch lengths prior to analysis by gene- and species-tree reconciliation in MAPS^116^.

Prior to running gene tree simulations, ultra-metric species trees containing taxa from each OrthoFinder run were generated using the R package ‘ape’^117^. Node ages were calibrated using maximum and minimum ages from Testo *et al.* ^30^. Next, estimates of background rates of gene duplication and loss across the tree were obtained using the R ‘WGDgc’ package^118^. To select nodes for positive simulations we initially utilized results from Ks and synteny analyses. In *Diphasiastrum complanatum* we observed a single peak in the Ks plot as well as a 1:1 relationship among collinear, intragenomic gene pairs. While we likewise observed a single peak in our Ks plot of collinear *Huperzia* genes, the extremely low orthologous divergence within Lycopodiaceae suggests that we may not be able to discern multiple peaks following the divergence of these two taxa. When combined with the overall 2:4 syntenic depth suggested by the results of MCSCAN we thus hypothesized two WGD events in *H. asiatica* for the one observed in *D. complanatum*. We therefore selected the node with the highest proportion of subtrees exhibiting a shared duplication in *D. complanatum* (N4) and the two nodes (N3 and N4) with the highest proportion of duplicated subtrees in *H. asiatica*. Despite N2 exhibiting slightly elevated retention of duplicated subtrees in MAPs analyses of both species, we did not carry out positive simulations based on (i) the expected total number of duplications from Ks and synteny analyses, (ii) the relatively low proportion of duplicated subtrees at N2 relative to other duplicated nodes, and (iii) the likelihood that duplications toward the tips of trees might be the result of shared tandem duplications and gene family expansions in closely related taxa or (in the case of *H. asiatica*) the result of subtrees duplicated at N3 where one paralog had been lost in either *H. asiatica* subA or subB. Finally, null and positive simulations were run using the ‘simulateGeneTrees.3.0.pl’ script included in MAPS. For *Diphasiastrum*, a single WGD was simulated at the last common ancestor (LCA) of *D. complanatum* and *Dendrolycopodium obscurum* with a retention rate of 0.4. For positive simulations with *Huperzia asiatica* as focal species, WGDs were simulated at the LCA of *H. asiatica* subA and *H. asiatica* subB as well as the LCA of each subgenome and *H. javanica*. MAPS was run on simulated gene trees with retention rates of 0.4 and 0.3, respectively. Finally, comparisons of the experimental and simulated results were assessed for significance using Fisher’s exact test in R.

### Substitution rate

Substitution rates across lycophytes were estimated with protein coding genes from *Diphasiastrum complanatum*, *Huperzia asiatica*, and transcriptomic data from *Phyloglossum drummondii*^9^, *H. javanica*^95^, *H. lucidula*^9^, *H. miyoshiana*^9^, *Phlegmariurus myrsinites*^9^, *P. squarrosa*^9^, *Dendrolycopodium obscurum*^9^, *Spinulum annotinum*^9^, *D. digitatum*^9^, *Lycopodiella appressa*^9^, *Selaginella moellendorffii*^10^, *S. tamariscina*^11^, *S. lepidophylla*^12^, *S. kraussiana*^11^, *Isoetes echinospora*^119^, *I. tegetiformans*^9^, *I. drummondii*^120^, *I. sinensis*^121^, and *I. taiwanensis*^13^. OrthoFinder^96^ was run to identify orthogroups and resulting orthogroups were filtered with a custom python script to remove taxa represented by more than one sequence, likely due to gene duplication. Downstream analysis was further restricted to orthogroups containing at least one species from each major clade (Lycopodiaceae, Selaginellaceae, and Isoetaceae) and with a minimum of 13 out of 21 taxa. Orthogroups were aligned in PRANK^122^ using the codon model with default settings prior to detecting shifts in substitution rate using PAML^97^. In order to produce the input phylogeny for baseml, 241 single copy gene alignments from OrthoFinder were used to produce gene trees using IQ-TREE^123^. Gene trees from IQ-TREE were subsequently used as input for ASTRAL^124^ to produce the final species phylogeny. The significance of rate change was inferred by a likelihood ratio test between two baseml models. The first model used a global clock with a single rate, and the second model used a local clock with 3 different rates (one for each lycophyte order).

## Data availability

The raw data of genome and transcriptome sequencing of *H. asiatica* have been deposited to the Genome Sequence Archive at the National Genomics Data Center (NGDC) under BioProject No. PRJCA013778. The genome assemblies and annotations of *H. asiatica* and all the chloroplast genomes assembled are available at figshare platform (https://figshare.com/projects/Huperzia_asiatica_genome/169145). The raw data of genome and transcriptome sequencing of *D. complanatum* have been deposited in the NCBI SRA under BioProject No. PRJNA914350. The genome assemblies and annotations of *D. complanatum* can be found in Phytozome (https://phytozome-next.jgi.doe.gov/info/Dcomplanatum_v3_1).

## Code availability

All custom codes are available for research purposes from the corresponding authors upon request.

## Acknowledgements

This study was supported by National Key Research and Development Program of China (Grant No. 2020YFA0907900), National Natural Science Foundation of China (Grant No. 32070242), Shenzhen Science and Technology Program (Grant No. KQTD2016113010482651), Special funds for Science Technology Innovation and Industrial Development of Shenzhen Dapeng New District (Grant Nos. RC201901-05 and PT201901-19), China Postdoctoral Science Foundation (Grant No. 2020M672904), Basic and Applied Basic Research Fund of Guangdong (Grant No. 2020A1515110912), and Science, Technology and Innovation Commission of Shenzhen Municipality of China (ZDSYS 20200811142605017). The work (proposal: 10.46936/10.25585/60001405) also conducted by the U.S. Department of Energy Joint Genome Institute (https://ror.org/04xm1d337), a DOE Office of Science User Facility, is supported by the Office of Science of the U.S. Department of Energy operated under Contract No. DE-AC02-05CH11231.

## Author contributions

L.W. and F.-W.L. conceived the project. C.L., L.-Y.K., X.C., B.N., J. Ji., X.H., S. Sun., and G.D. undertook the sampling and experimental work. C.L., J. Jenkins, M.W., S. Shu., C.P., K.B., S.R., J.G., J.S., and J.H.L.-M. conducted the genome assemblies and annotations. C.L., D.W., X.L., D.P., Z.H., W.H., and C.S. performed data analyses. C.L., D.W., F.-W.L., and L.W. synthesized and wrote the manuscript.

