## Supplementary text and figure1-26 for "Extraordinary preservation of gene collinearity over three hundred million years revealed in homosporous lycophytes"

### SUPPLEMENTARY INFORMATION

- **Supplementary Text**
  - Subgenome phasing for the allotetraploid *Huperzia asiatica*
  - Evolution of gene families and transcription factors
- **Supplementary Methods**
  - Transcription-associated protein (TAPs) identification
  - Identification of *YABBY* genes
  - Phylogeny and gene family evolution
  - Identification of TMO5 and LHW genes and evolution
- **Supplementary Fig. 1.** K-mer (23-mer) distribution and estimation of genome size of *H. asiatica* (a) and *D. complanatum* (b).
- **Supplementary Fig. 2.** Hi-C interaction maps for *D. complanatum* (a) and (b) *H. asiatica* genomes.
- **Supplementary Fig. 3.** Principal component analysis (PCA) (a) and heatmap clustering (b) of subgenome-specific 15-mers in *H. asiatica*.
- **Supplementary Fig. 4.** Subgenome-specific k-mer and repeat characteristics in *H. asiatica*.
- **Supplementary Fig. 5.** Comparisons of chromosome length (a), gene number (b), and GC content (c) between *H. asiatica* subgenomes.
- **Supplementary Fig. 6.** Syntenic relationship between *H. asiatica* subgenomes.
- **Supplementary Fig. 7.** Analysis of genes involved in two large-scale chromosomal rearrangements in *H. asiatica.*
- **Supplementary Fig. 8.** Comparison of orphan gene number between *H. asiatica* subgenomes.
- **Supplementary Fig. 9.** Homolog expression bias (HEB) across the six tissues in the expression atlas of *H. asiatica*.
- **Supplementary Fig. 10.** Comparisons for absolute value of $\log_2$FoldChange of homoeolog expression bias (HEB; cutoffs: '*P*-value < 0.05 and | $\log_2$FoldChange | > 1') gene pairs between *H. asiatica* subgenomes in six different tissue types.

- **Supplementary Fig. 11.** Different LTR coverage in the flanking regions between the dominant and the suppressed member for each homolog expression bias gene pairs (cutoffs: ‘*P*-value < 0.05 and | $\log_2$FoldChange | > 1’) of *H. asiatica* in different tissue types.
- **Supplementary Fig. 12.** Plots of Inter-genomic synteny and relative syntenic depth between *H. asiatica* and *D. complanatum* genomes.
- **Supplementary Fig. 13.** Plots of intra-genomic synteny in *H. asiatica* and *D. complanatum* genomes.
- **Supplementary Fig. 14.** Diagram of hypothetical syntenic relationships following multiple rounds of WGD and reciprocal gene fractionation.
- **Supplementary Fig. 15.** An in-depth example of collinear blocks exhibiting 2:4 syntenic depth ratio between *D. complanatum* and *H. asiatica* subA.
- **Supplementary Fig. 16.** Collinear genes plotted on scaffolds showing multiple examples of 2:4 syntenic depth between *D. complanatum* and *H. asiatica* subA.
- **Supplementary Fig. 17.** Collinear genes plotted on scaffolds showing multiple examples of 2:4 syntenic depth between *D. complanatum* and *H. asiatica* subB.
- **Supplementary Fig. 18.** Ks plots for species in Lycopodioideae and Lycopodielloideae.
- **Supplementary Fig. 19.** Ks plots for species in Huperzioideae and Lycopodioideae.
- **Supplementary Fig. 20.** Summary of MAPs analysis focused on *H. asiatica* subB.
- **Supplementary Fig. 21.** Gene family evolution of representative land plant lineages.
- **Supplementary Fig. 22.** GO terms in the ‘Biological Process’ category that are associated with the expanded gene families in the common ancestor of embryophytes, tracheophytes, and euphyllophytes.
- **Supplementary Fig. 23.** The gene number of representative transcription factors (TFs) families in land plant lineages.
- **Supplementary Fig. 24.** Phylogenetic analysis of TMO5 proteins in *H. asiatica*, *D. complanatum* and representative species among green plants.

- **Supplementary Fig. 25.** Phylogenetic analysis of LHW proteins in *H. asiatica*, *D. complanatum* and representative species among green plants.
- **Supplementary Fig. 26.** Phylogenetic profiling of representative species among green plants.
- **Supplementary Table 1.** Assembly and annotation statistics of *Huperzia asiatica* and *Diphasiastrum complanatum* genomes.
- **Supplementary Table 2.** Mapping rates of genomic and transcriptomic Illumina reads.
- **Supplementary Table 3.** Statistics of transposable elements (TEs) in *Huperzia asiatica*, *Diphasiastrum complanatum* and other lycophyte genomes.
- **Supplementary Table 4.** BUSCO assessments of *Huperzia asiatica*, *Diphasiastrum complanatum* and other lycophyte genomes and proteomes.
- **Supplementary Table 5.** Gene length statistics of *Huperzia asiatica*, *Diphasiastrum complanatum* and other representative species in green plants.
- **Supplementary Table 6.** Comparison of LTR insertion in intron between homosporous and heterosporous lycophytes.
- **Supplementary Table 7.** Number and proportion of transcription associated proteins (TAPs) in *Huperzia asiatica*, *Diphasiastrum complanatum* and other selected plant species.
- **Supplementary Table 8.** The 18,264 homolog gene pairs with the corresponding positions between *Huperzia asiatica* subgenomes.
- **Supplementary Table 9.** Whole genome sequencing information of *Huperzia* species and *Diphasiastrum complanatum*.
- **Supplementary Table 10.** RNA-Seq information of *Huperzia* species and *Diphasiastrum complanatum*.
- **Supplementary Table 11.** Chromosome length and GC content in *Huperzia asiatica* genome.
- **Supplementary Table 12.** Significantly enriched GO terms for the expanded gene families of *Huperzia asiatica* (p.adjust < 0.01).
- **Supplementary Table 13.** Significantly enriched GO terms for the expanded gene families of *Diphasiastrum complanatum* (p.adjust < 0.01).

- **Supplementary Table 14.** GO terms in the 'Biological Process' category that are associated with the expanded gene families in the common ancestor of embryophytes.
- **Supplementary Table 15.** GO terms in the 'Biological Process' category that are associated with the expanded gene families in the common ancestor of tracheophytes.
- **Supplementary Table 16.** GO terms in the 'Biological Process' category that are associated with the expanded gene families in the common ancestor of tracheophytes.

### Supplementary Text:

**Subgenome phasing for the allotetraploid *Huperzia asiatica*.** To phase the allotetraploid genome, we used k-mer sequences, which can reflect the genomic features (e.g., repeat types and frequencies) that are characteristic of each subgenome. Similar approach has been used in several allotetraploid species, including *Panicum virgatum*, *Miscanthus sinensis* and *Eragrostis teff*[1-3]. We found 7,579 15-mers whose pairwise enrichment pattern consistently partitions the homoeologous chromosome pairs between distinct A and B subgenomes (hereafter referred "subA" and "subB") (Supplementary Figs. 3,4a-c). We also identified LTR-RTs that are enriched in one member of each homeologous chromosome pair between two subgenomes (Supplementary Fig. 4d-f), which confirms the k-mer-based subgenome phasing results. *H. asiatica* subA and subB are consisted of 70 and 68 chromosomes with average chromosome size of 50.9 Mb and 62.7 Mb, respectively (Supplementary Fig. 5a and Supplementary Table 11). *H. asiatica* subB harbours more genes and TEs than the subA (Supplementary Fig. 5b and Supplementary Table 3), while the GC content in subA is higher than that in subB (Supplementary Fig. 5c and Supplementary Table 11).

**Evolution of gene families and transcription factors.** Gene families and ortholog sets were circumscribed for gene models characterized for *H. asiatica*, *D. complanatum* and other 21 published genomes spanning major lineages of land plants and streptophyte algae (Supplementary Fig. 21). Phylogenetic analysis of low-copy orthologous groups confirmed Lycopodiales is sister to a clade consisting of Selaginellales and Isoetales, and the divergence time of Lycopodielloideae+Lycopodioideae (represented by *D. complanatum*) and the Huperzioideae was estimated to be 335.7 MYA (95% confidence interval: 323.1-371.0 MYA). Gene family evolution analysis identified 211 families expanded and 31 families contracted along the branch leading to Lycopodiaceae. Gene Ontology (GO) analysis of the 301 and 268 expanded gene families in *H. asiatica* and *D. complanatum* resulted in 700 and 381 significantly enriched GO terms, respectively (Supplementary Tables 12 and 13). These include those involved in the responses to stimulus (e.g., drought stress, ultraviolet light, and wounding) and the biosynthesis of plant hormones (e.g., brassinosteroids, gibberellins, and jasmonates) and secondary metabolites (e.g., terpenoids and phenylpropanoids. We also investigated changes in gene family content and size associated with plant terrestrialization and subsequent environmental adaptation. Bursts of gene family expansion were evident on branches leading to the common ancestors of embryophytes (288), tracheophytes (320) and euphyllophytes (133). Most of the expanded gene families on these ancestor nodes are annotated with functions related to multicellularity and increasing anatomical complexity (Supplementary Fig. 22 and Supplementary Tables 14-16). Other substantially expanded gene families are associated with responses to stresses, endogenous stimulus and hormones. These results suggest that the transition to novel terrestrial environments was accompanied by the sustained expansion of gene families involved in organ development and stress adaptability.

We found that 5.2% and 6.1% of the proteomes of *H. asiactica* and *D. complanatum* genomes were annotated as transcription-associated proteins (TAPs), which include transcription factors (TFs) and transcription regulators (TRs) (Supplementary Table 7). *H. asiactica* and *D. complanatum* genomes contain a higher proportion of TAPs compared to bryophytes. The evolution of *YABBY*, a TF gene family that plays a key role in specifying leaf adaxial-abaxial polarity in seed plants[4-6], is particularly noteworthy. One *YABBY* ortholog was detected in *D. complanatum*, but none in *H. asiatica* despite the previous report of a single *YABBY* homolog in *H. selago* transcriptome data[7]. We then searched transcriptomes of *H. miyoshiana*, *H. lucidula*, *H. javanica*, and *H. serrata*, and no *YABBY* homolog was identified. These results supported that *YABBY* may have been lost multiple times in *Huperzia* species, although the incompleteness in transcriptome data might also contribute to the spotty distribution. Moreover, taking account of previous reports that *YABBY* was absent in genomes of mosses[8], liverworts[9], *Isoetes*[10], *Selaginella*[11], and ferns[12,13], but present in hornworts[14], the repeated loss of *YABBY* appears to be not necessary for abaxial/adaxial polarity among seed-free lineages. The dynamic evolution of *YABBY* is perplexing and warrants further research. The bHLH, NAC, MYB and WRKY TF families, all of which are involved in regulation of organ development[9,14,15], exhibit sustained expansion in vascular plants (Supplementary Fig. 23). For example, the heterodimeric TMO5/LHW bHLH TF complex is rate-limiting for vascular cell proliferation in vascular plants[16,17]. Although the orthologs of TMO5 and LHW can be found in bryophyte lineages (Supplementary Figs. 24 and 25), the generation of the critical function for the complex is evolved in vascular plants[16]. Moreover, we found that the copy numbers of TMO5 and LHW TFs expanded accompanied by WGDs, which occurred in different lineages of vascular plants (Supplementary Fig. 26).

### Supplementary Methods:

**Transcription-associated protein (TAPs) identification.** Transcription-associated proteins (TAPs) include transcription factors (TFs) that bind in a sequence-specific manner to *cis*-regulatory DNA elements, and transcription regulators (TRs) that act through protein-protein interaction or chromatin modification. To identify and compare TAPs among land plants, iTAK[18] was utilized to identify TFs and TRs from protein sequences, and the individual TFs and TRs were classified into different gene families. The known plant TFs and TRs present in the iTAK database (http://itak.feilab.net/cgi-bin/itak/online_itak.cgi) were used as a reference. To avoid missing family members with low identity value, the protein sequences of each putative TF or TR from *Arabidopsis thaliana* were used as queries in a BLASTP search of the lycophyte species, including *Huperzia asiatica*, *H. miyoshiana*, *H. lucidula*[19], *H. javanica*[20], *H. serrata*, *Diphasiastrum complanatum*, and *Isoetes taiwanensis*[10], protein datasets with cutoff '-e-value $1.0\times10^{-10}$'. The conserved domains in all the putative TF/TR identified were examined with CD-Search[21].

**Identification of *YABBY* genes.** We initially extracted *YABBY* genes in our iTAK results. To confirm the result, *YABBY* genes were identified from Orthofinder results, based on our assembled genomes of *Huperzia asiatica* and *Diphasiastrum complanatum*, OneKP[19] transcriptome of *H. lucidula* and our assembled transcriptomes of *H. serrata*, *H. miyoshiana*, and *H. javanica*, as well as the genomes from other 21 representative species among green plants, if they belonged to any group containing genes identified in the previous step or a group having known *YABBY* genes from *Arabidopsis thaliana*, *Anthoceros agrestis*, and *Amborella trichopoda.* Conserved domains were also used as search queries against the predicted proteome using HMMER (https://www.ebi.ac.uk/Tools/hmmer/).

**Phylogeny and gene family evolution.** *H. asiatica*, *D. complanatum* and nineteen other species representing major lineages in Viridiplantae, including *Klebsormidium nitens*[22], *Chara braunii*[23], *Spirogloea muscicola*[15], *Mesotaenium endlicherianum*[15], *Penium margaritaceum*[24], *Anthoceros agrestis* Bonn[14], *Physcomitrium patens*[8], *Marchantia polymorpha*[9], *Isoetes taiwanensis*[10], *Selaginella moellendorffii*[11], *Selaginella lepidophylla*[25], *Azolla filiculoides*[13], *Salvinia cucullata*[13], *Adiantum capillus-veneris*[26], *Ceratopteris richardii*[27], *Alsophila spinulosa*[12], *Ginkgo biloba*[28], *Picea abies*[29], *Amborella trichopoda*[30], *Oryza sativa*[31] and *Arabidopsis thaliana*[32], were used for comparative genomic analysis. OrthoFinder[33] was applied to cluster orthologous groups among these species with parameters '-M msa -S diamond'. Here, 771 low-copy orthologous groups (i.e. at most three gene copies of each orthologous group for each species) were selected for the construction of the phylogenetic tree. Protein sequences were filtered by choosing the longest isoform to represent protein for each species in orthologous groups. Each orthologous group was aligned separately using multiple sequence alignment by MAFFT[34] and poor quality alignments were filtered out using trimAl[35]. The aligned sequences were concatenated into one super-sequence for each species and then the maximum likelihood (ML) tree was inferred using RAxML[36] with parameters '-f a -N 1000 -m PROTGAMMAJTT -x 123456 -p 123456' and *Klebsormidium nitens* as outgroup. For the coalescence-based phylogeny, 771 gene trees were separately constructed by RAxML with 500 bootstrap replicates, and then ASTRAL[37] was implemented to summarize the gene trees. We obtained the identical topology through concatenation and coalescence methods. Molecular dating was carried out by MCMCTree program in PAML[38] with the following constraints: (i) 637-875 MYA for Streptophyta[39], (ii) 482-515 MYA for Embryophyta[39], (iii) 430-452 MYA for Tracheophyta[39], (iv) 402-436 MYA for Euphyllophyta[39], (v) 393-431 MYA for Lycopodiophyta[39], (vi) 330-365 MYA for Spermatophyta[39], (vii) 281-288 MYA for Salviniales + Cyatheales[40], (viii) 91-99 MYA for Salviniales[40] and (ix) 326-389 MYA for Lycopodiaceae[41]. The program discarded the first 5,00,000 iterations (parameter 'burnin'), and then it sampled every 50 iterations (parameter ‘sampfreq’) until it had gathered 1,000,000 samples (parameter ‘nsample’). In total, the MCMC ran for 50,500,000 (5,00,000 + 50 × 1,000,000) iterations. The resulting trace file of the MCMC program was evaluated using Tracer[42] to ensure convergence, and the effective sample size of all parameters should be confirmed to be at least 200. The gene family evolution was evaluated with CAFÉ[43] based on results from OrthoFinder. A cut-off *P*-value was calculated for each gene family, and families with *P* values lower than 0.05 were considered to experience a significantly accelerated rate of expansion or contraction.

**Identification of TMO5 and LHW genes and evolution.** We used BLASTP to search for TMO5 and LHW homoeologs in *H. asiatica*, *D. complanatum* and representative species among green plants using *Arabidopsis thaliana* TMO5 and LHW protein sequences as queries with cutoff ‘-e-value $1.0 \times 10^{-10}$’. HMMER was also used in conserved domain building based on *A. thaliana* TMO5 and LHW sequences and predicted protein searching. To explore the phylogeny of TMO5 and LHW in green plants, we selected TMO5 sequences from charophytes sequences and LHW sequences from bryophytes as the outgroups, respectively. The protein sequences were aligned with MAFFT[34], poor quality alignments were filtered out with trimAl[35], and the maximum likelihood (ML) trees of TMO5 and LHW were inferred using RAxML[36] with 500 bootstrap replicates.

### Supplementary Figures:

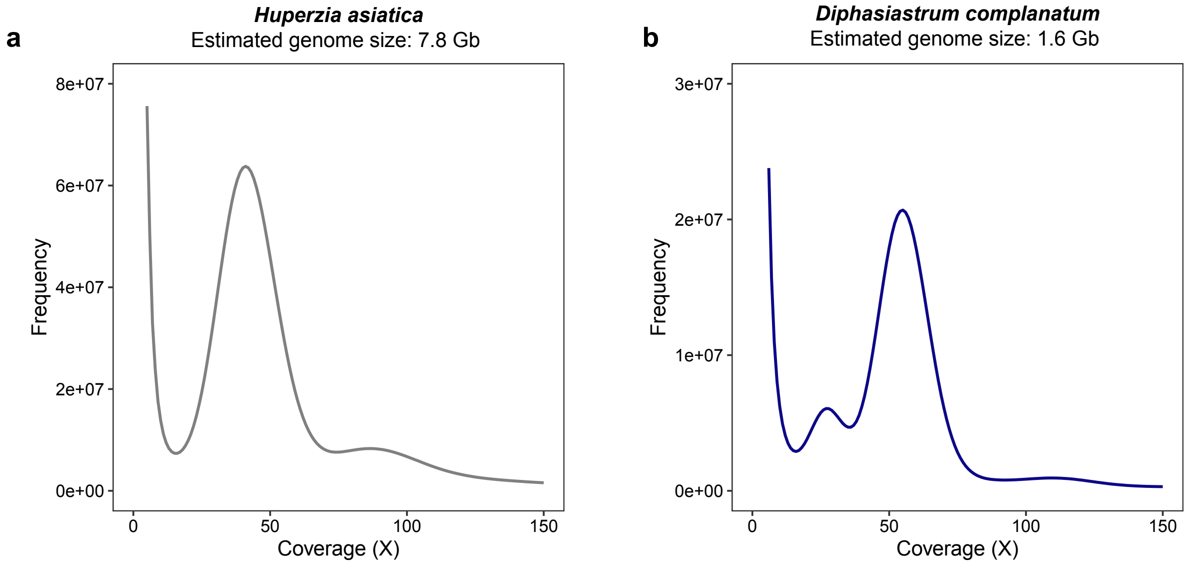


**Supplementary Fig. 1. K-mer (23-mer) distribution and estimation of genome size of *H. asiatica* (a) and *D. complanatum* (b).**

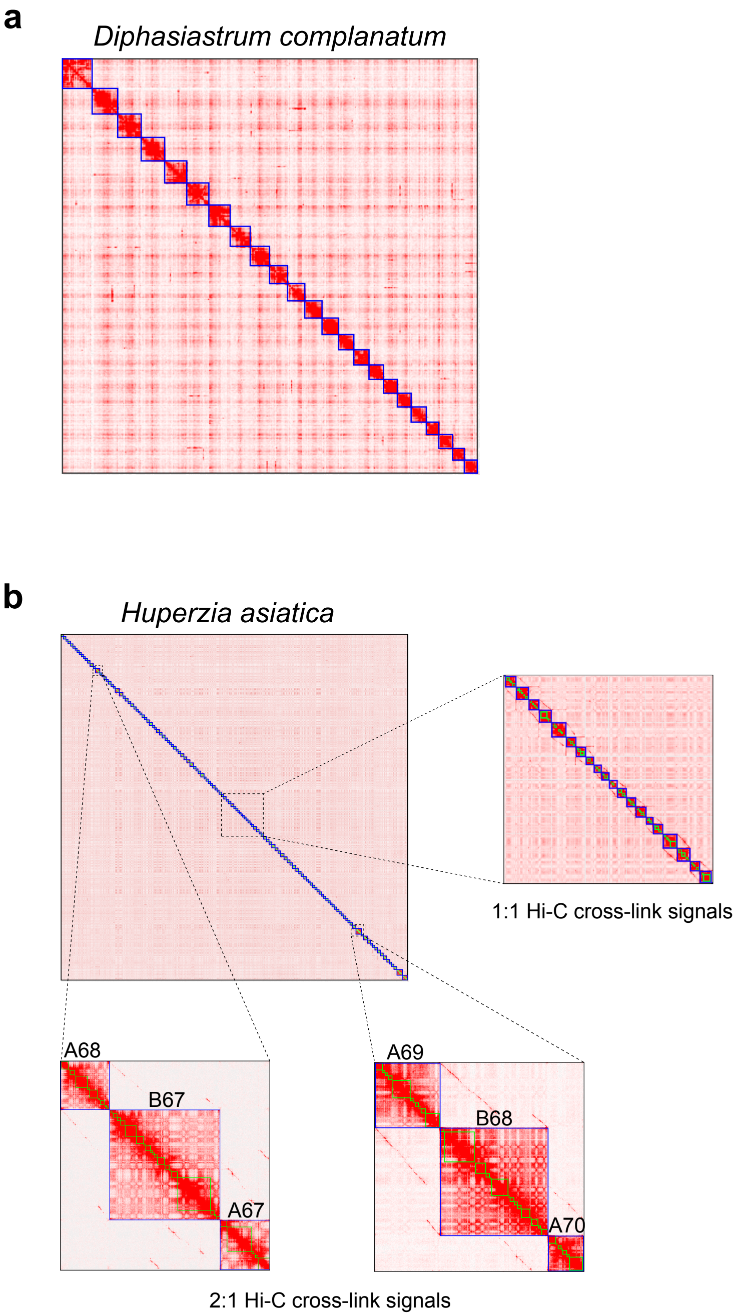


**Supplementary Fig. 2. Hi-C interaction maps for *D. complanatum* (a) and (b) *H. asiatica* genomes.** "1:1" or "2:1" Hi-C cross-link signals between chromosomes indicates the homoeolog chromosomes between *H. asiatica* subgenomes. Two large-scale "2:1" Hi-C cross-link signals, including "(A68+A67):B67" and "(A69+A70):B68", indicates either chromosome fusion or fission events between *H. asiatica* subgenomes.

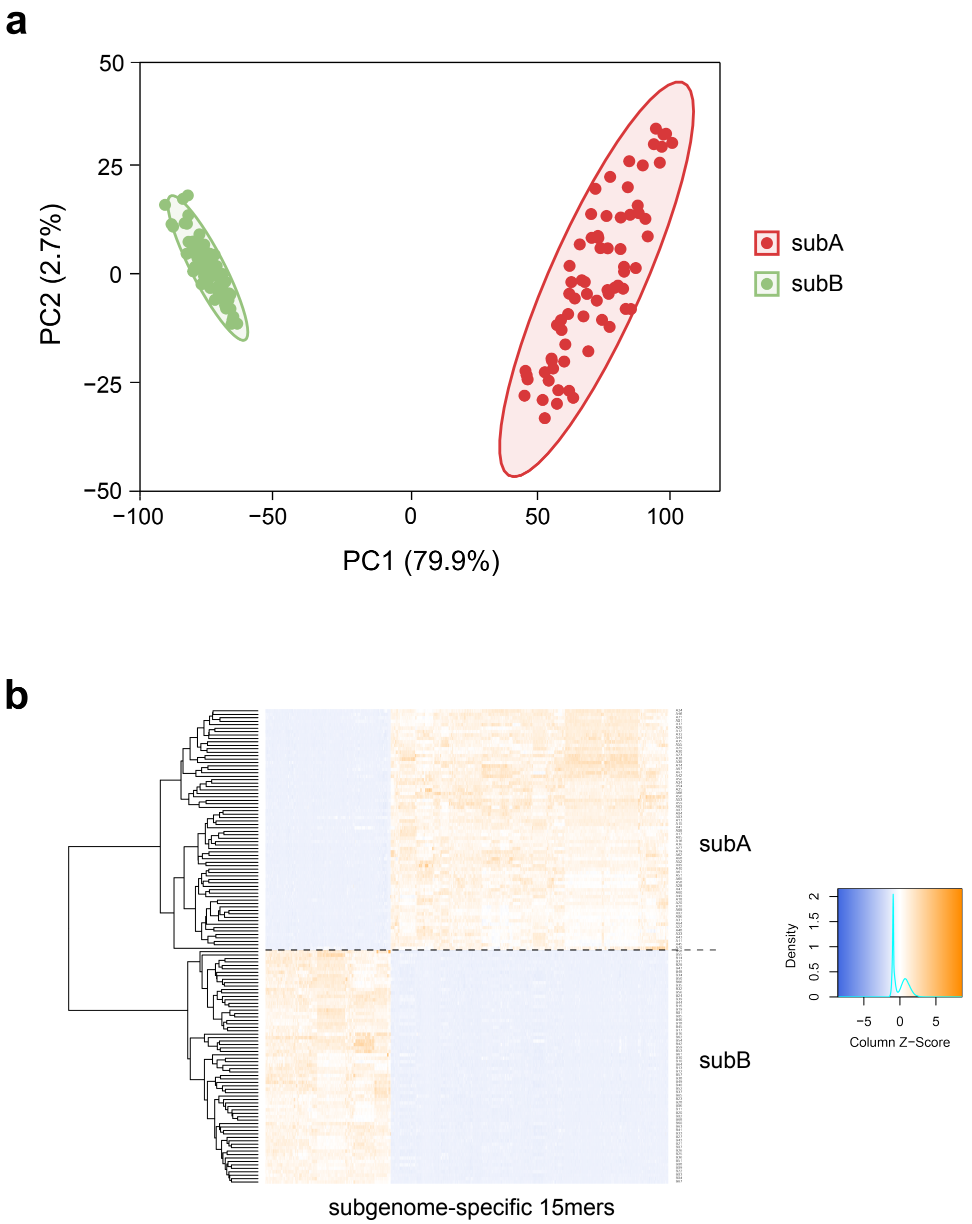


**Supplementary Fig. 3. Principal component analysis (PCA) (a) and heatmap clustering (b) of subgenome-specific 15-mers in *H. asiatica*.** The heatmap indicates the Z-scored relative abundance of 15-mers. The larger the Z-score, the higher the relative abundance of a 15-mer.

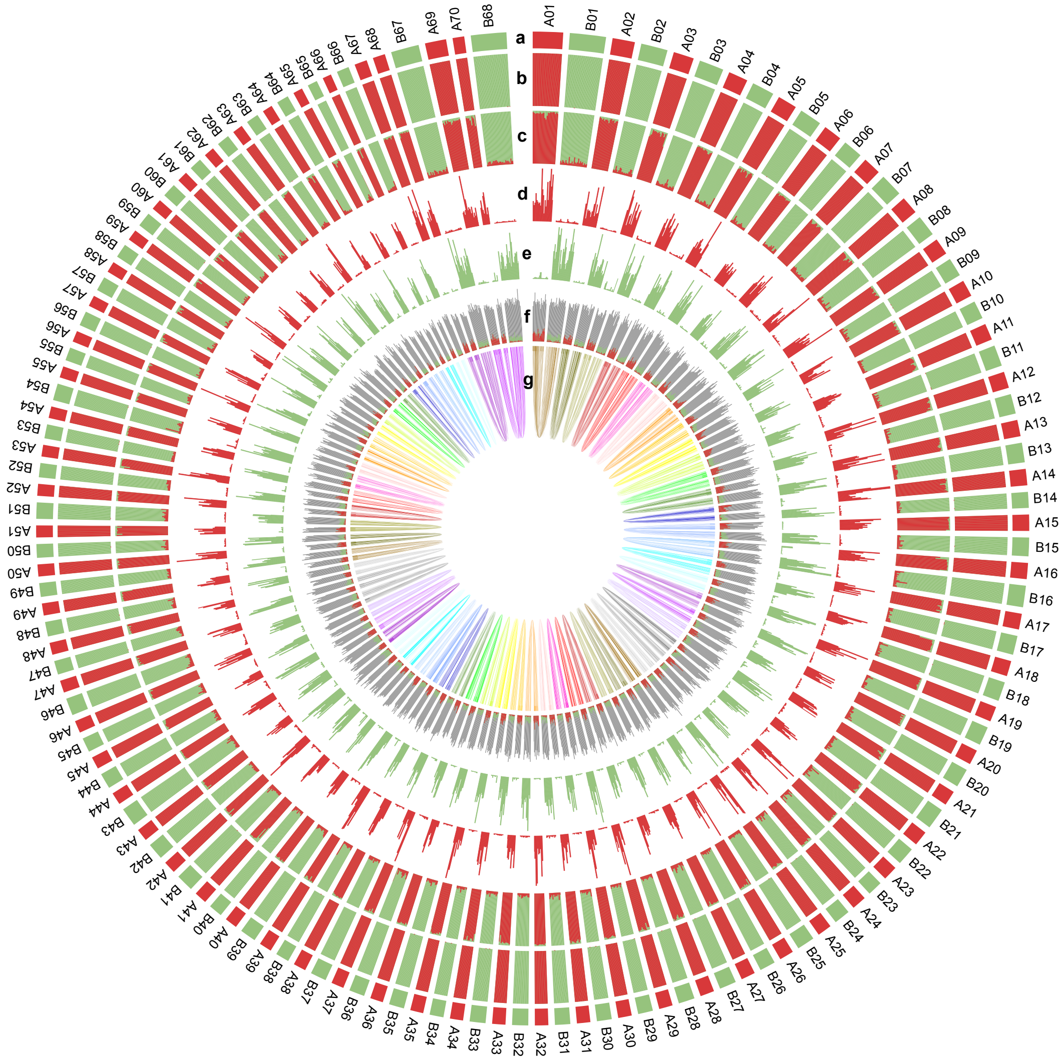


**Supplementary Fig. 4. Subgenome-specific k-mer and repeat characteristics in *H. asiatica*. a,** Subgenome assignments based on k-means algorithm. **b,** Significant enrichment of subgenome-specific 15-mers – the same color as the subgenome indicates significant enrichment for those subgenome-specific 15-mers. No evidence of recently large-scale homolog exchange was inferred by subgenome-specific 15-mers. **c,** Normalized proportion (relative) of subgenome-specific 15-mers. **d,e,** Count (absolute) of each subgenome-specific 15-mer set. **f,** Density of long terminal repeat retrotransposons (LTR-RTs) – if the color is consistent with the subgenome, it indicates that LTR-RTs are significantly enriched to those subgenome-specific 15-mers. Gray indicates nonspecific LTR-RTs. **g,** Homoeologous blocks. All statistics (**b,c,d,e,f,g**) are computed in sliding windows of 1 Mb. Red and green (**a,b,c,d,e,f**) indicate subA and subB of *H. asiatica*, respectively.

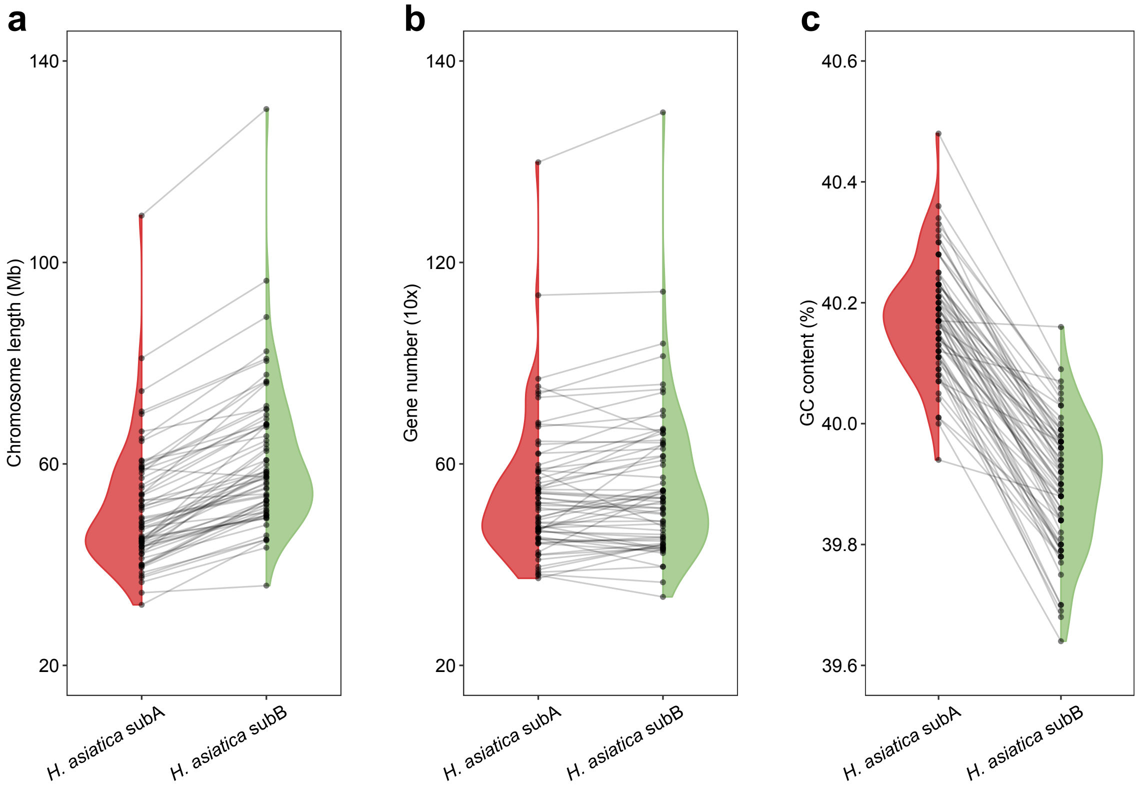


**Supplementary Fig. 5. Comparisons of chromosome length (a), gene number (b), and GC content (c) between *H. asiatica* subgenomes.** Points indicate chromosomes.

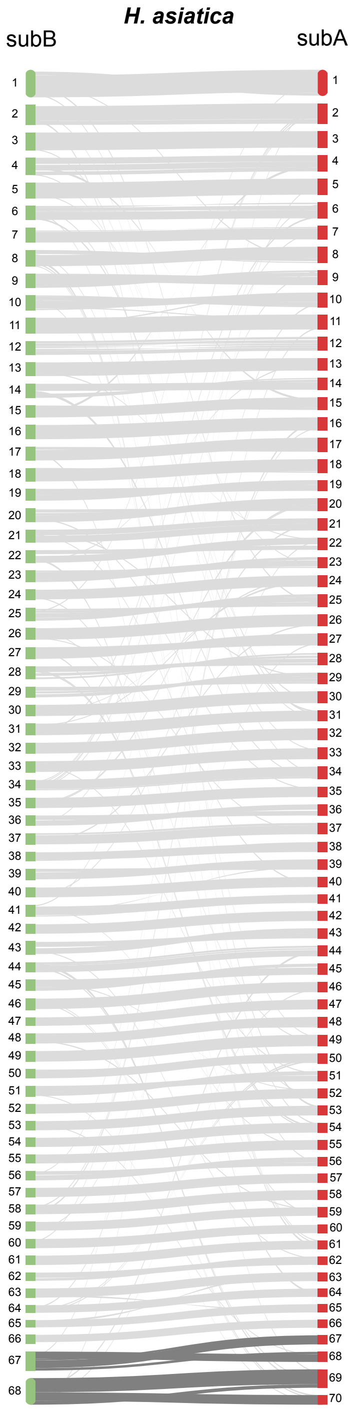


**Supplementary Fig. 6. Syntenic relationship between *H. asiatica* subgenomes.** Synteny blocks are connected by light gray lines and the inferred fusions or fissions are highlighted by dark gray lines.

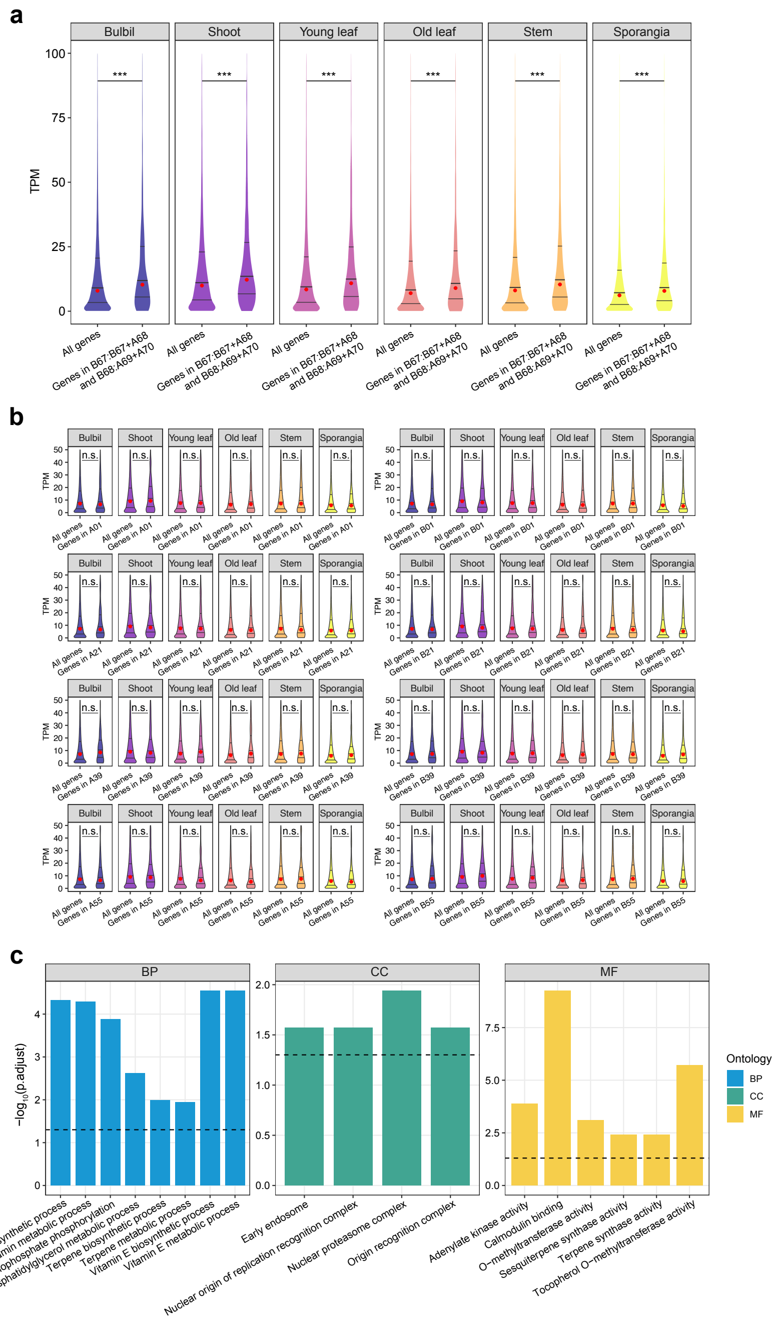


**Supplementary Fig. 7. Analysis of genes involved in two large-scale chromosomal rearrangements in *H. asiatica*. a,** Comparison of the expression levels of genes within the two chromosomal rearrangements and the genomic background. *P*-value estimated by Wilcoxon rank-sum test. '***' indicates *P*-value < $2.2 \times 10^{-16}$. **b,** Comparison of the expression levels of genes within the randomly selected four pairs of homoeologous chromosomes and the genomic background. *P*-value estimated by Wilcoxon rank-sum test. 'n.s' indicates no significant difference. **c,** Gene Ontology (GO) enrichment analysis for the genes involved in the two chromosomal rearrangements.

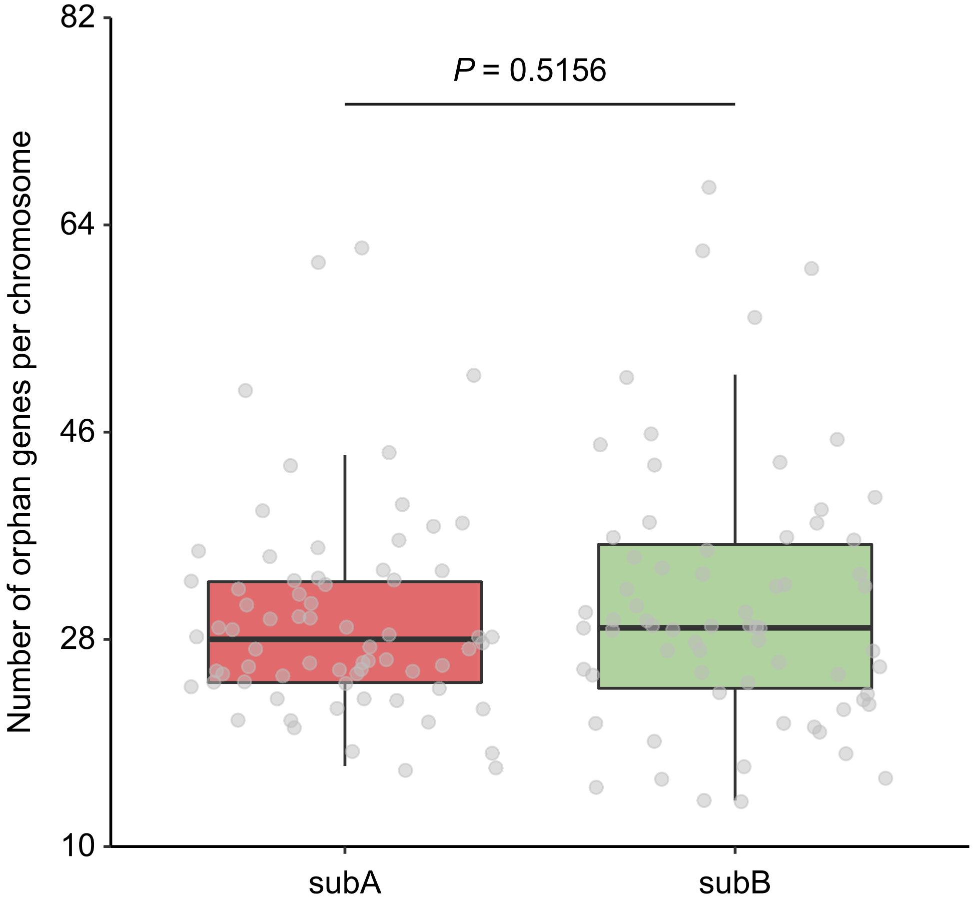


**Supplementary Fig. 8. Comparison of orphan gene number between *H. asiatica* subgenomes.** *P*-value was estimated by Wilcoxon rank-sum test. Orphan genes here are defined as those whose homoeologue was lost in the other subgenome.

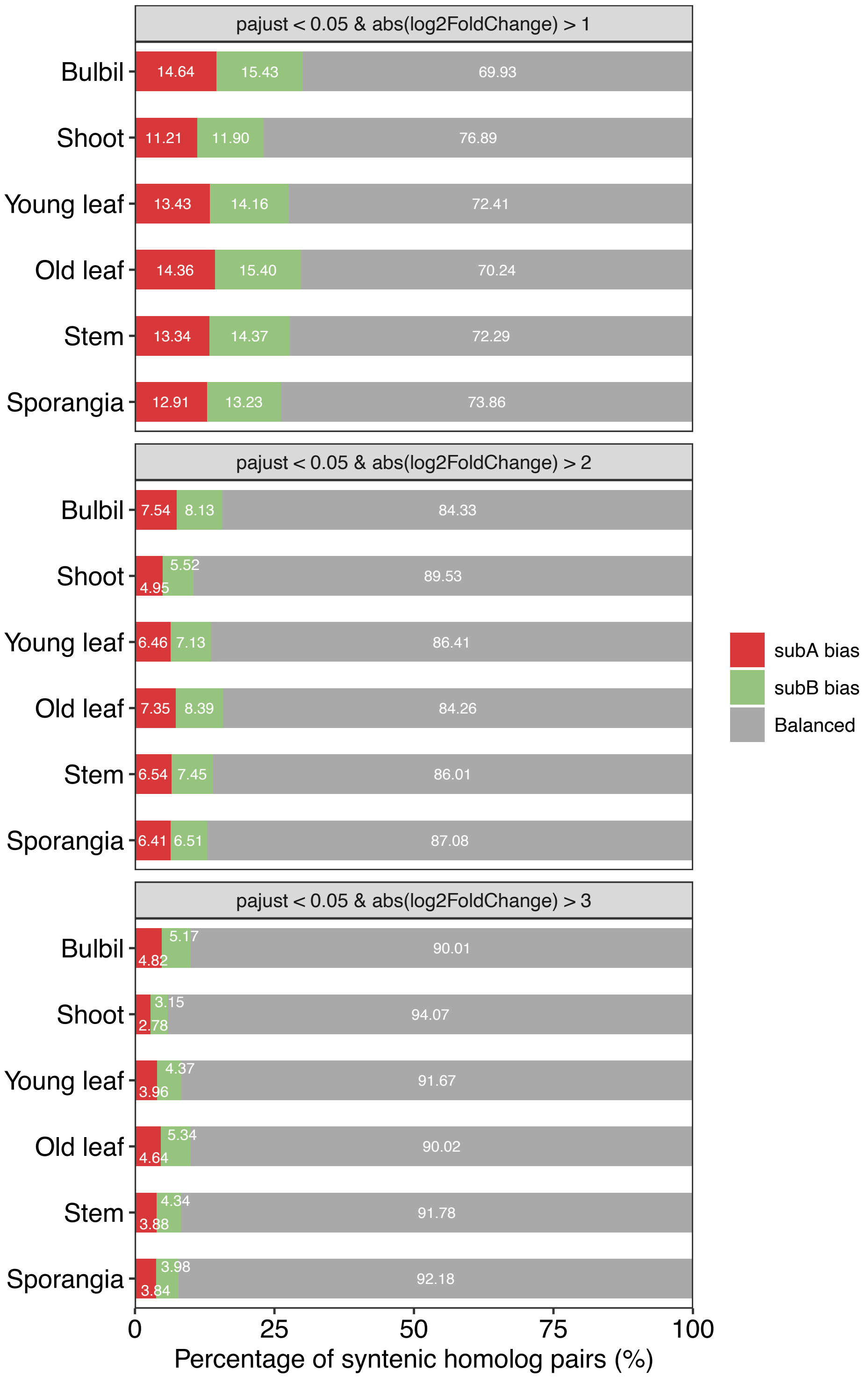


**Supplementary Fig. 9. Homoeolog expression bias (HEB) across the six tissues in the expression atlas of *H. asiatica*.** Gene pairs were classified as biased toward subA (red) or subB (green) or balanced with no statistically significant differential expression (gray).

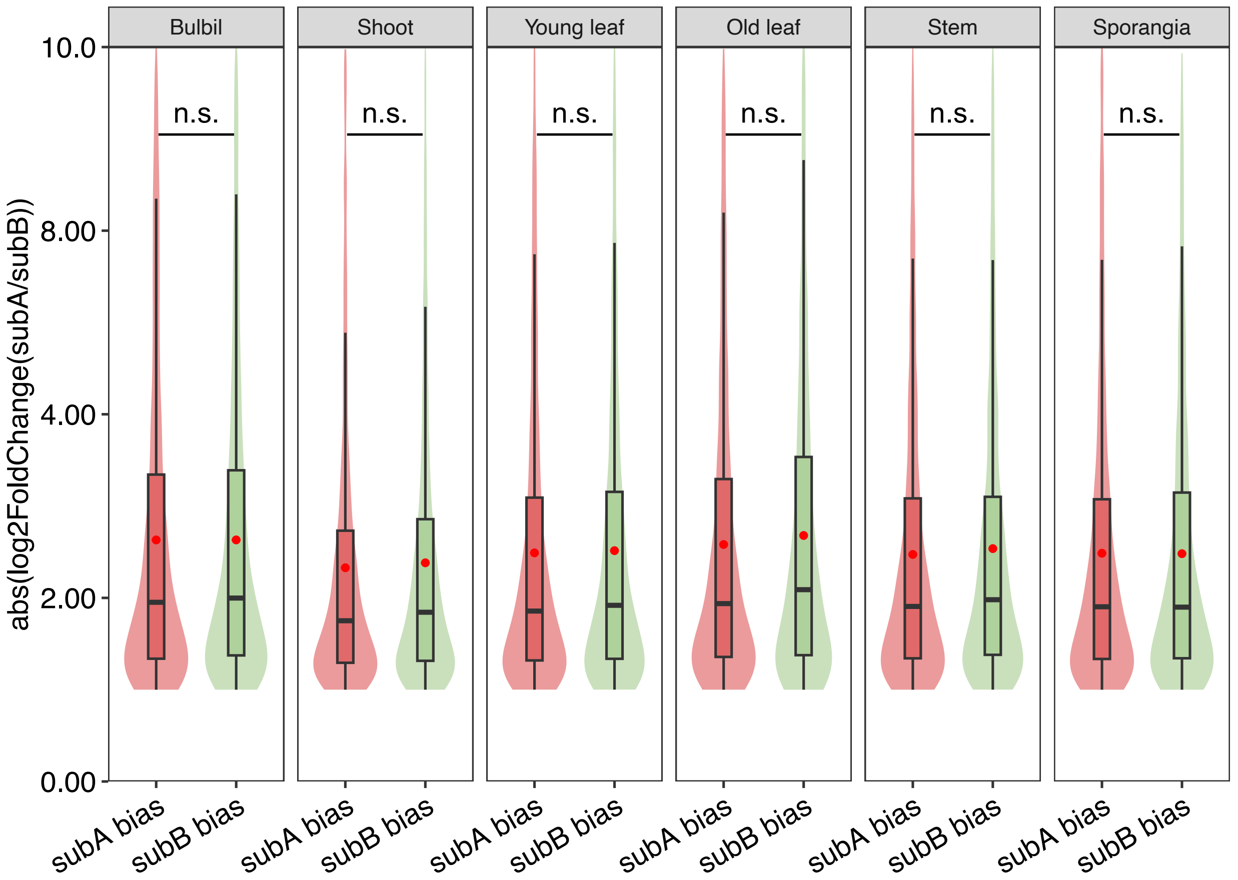


**Supplementary Fig. 10. Comparisons for absolute value of $\log_2$FoldChange of homoeolog expression bias (HEB; cutoffs: '*P*-value < 0.05 and | $\log_2$FoldChange | > 1') gene pairs between *H. asiatica* subgenomes in six different tissue types.** *P*-value was estimated by Wilcoxon rank-sum test. 'n.s' indicates no significant difference.

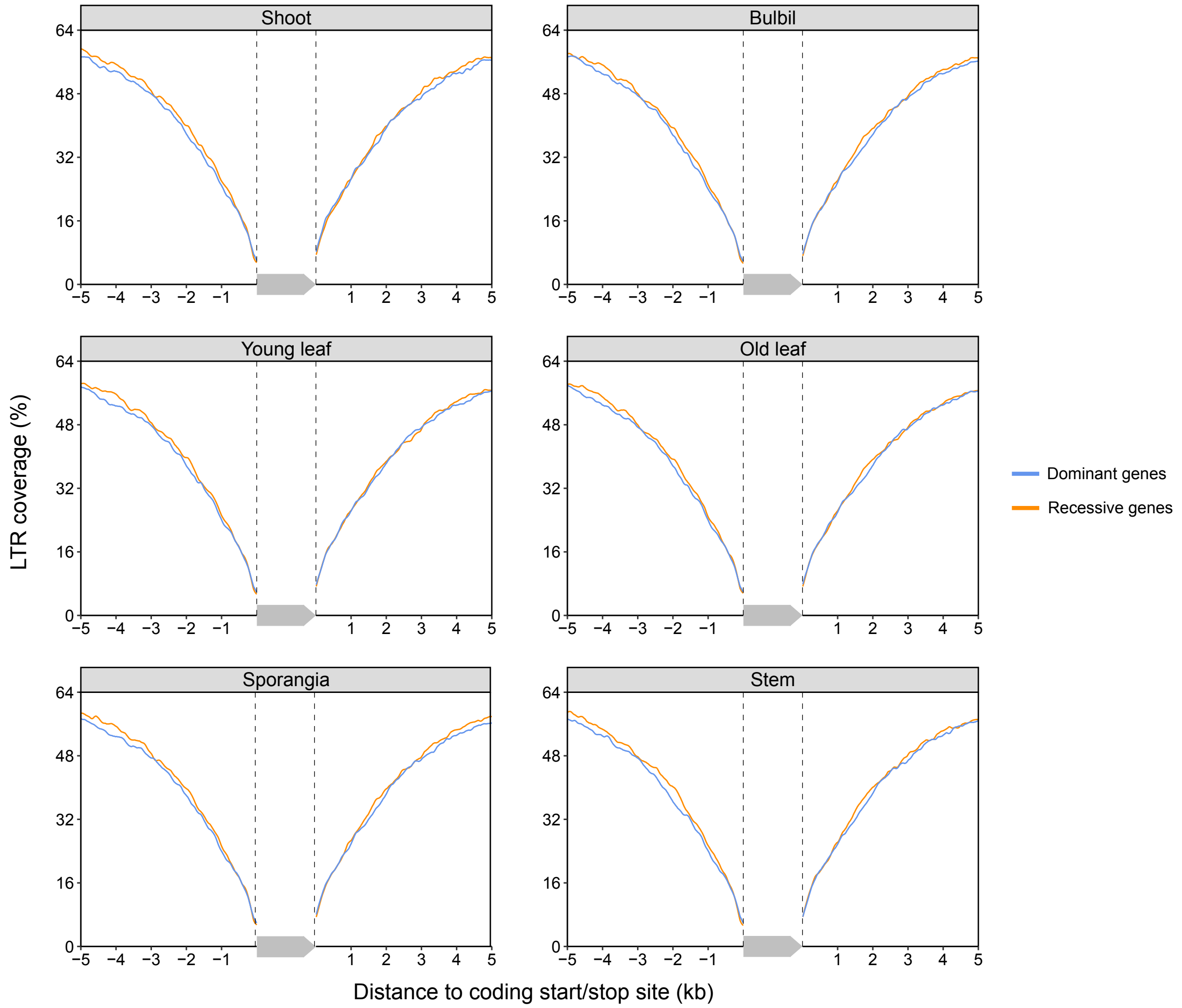


**Supplementary Fig. 11. Different LTR coverage in the flanking regions between the dominant and the suppressed member for each homolog expression bias gene pairs (cutoffs: '*P*-value < 0.05 and | $\log_2$FoldChange | > 1') of *H. asiatica* in different tissue types.** The two dashed lines and a gray arrow indicate the gene region and orientation, respectively.

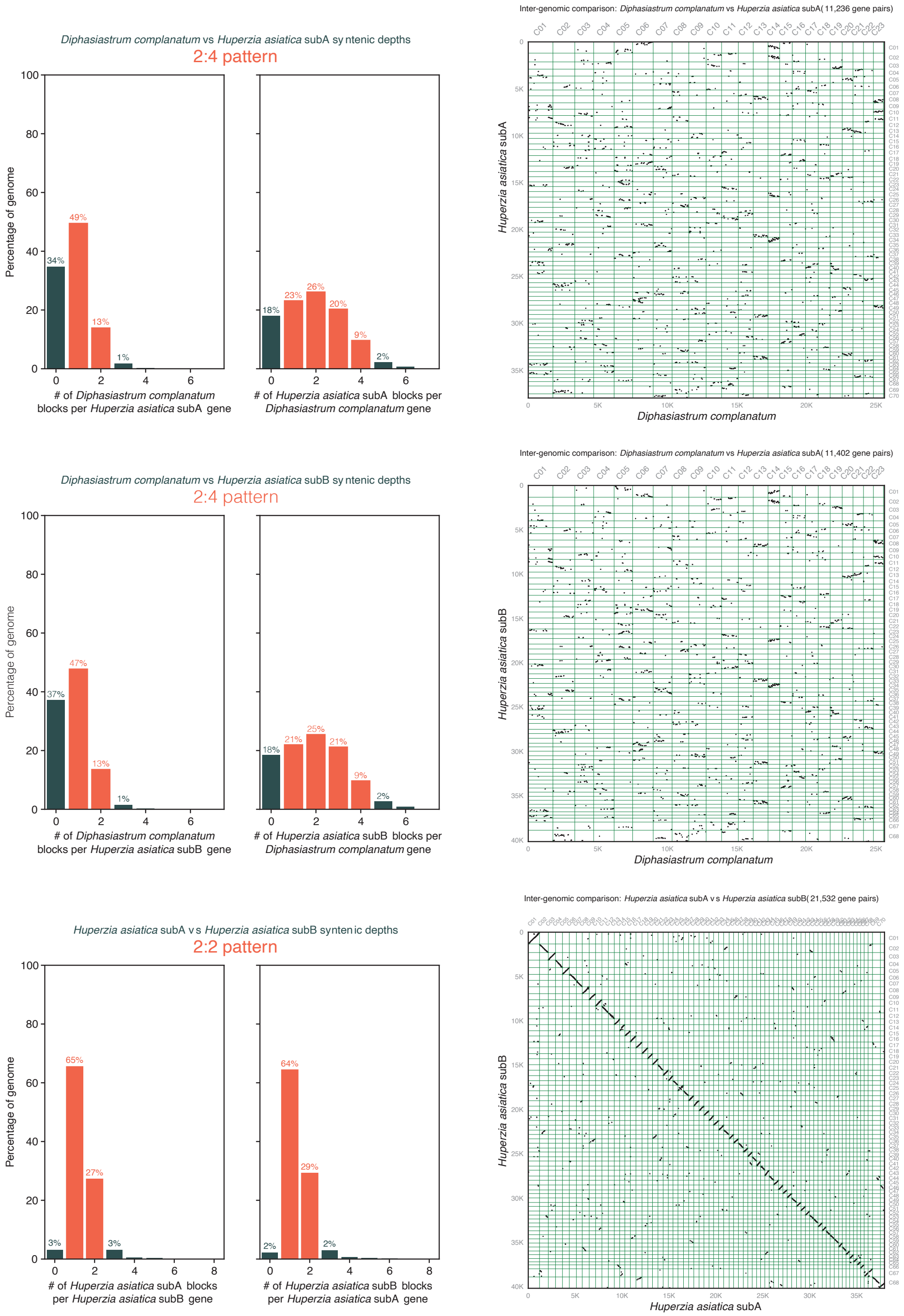


**Supplementary Fig. 12. Plots of Inter-genomic synteny and relative syntenic depth between *H. asiatica* and *D. complanatum* genomes.**

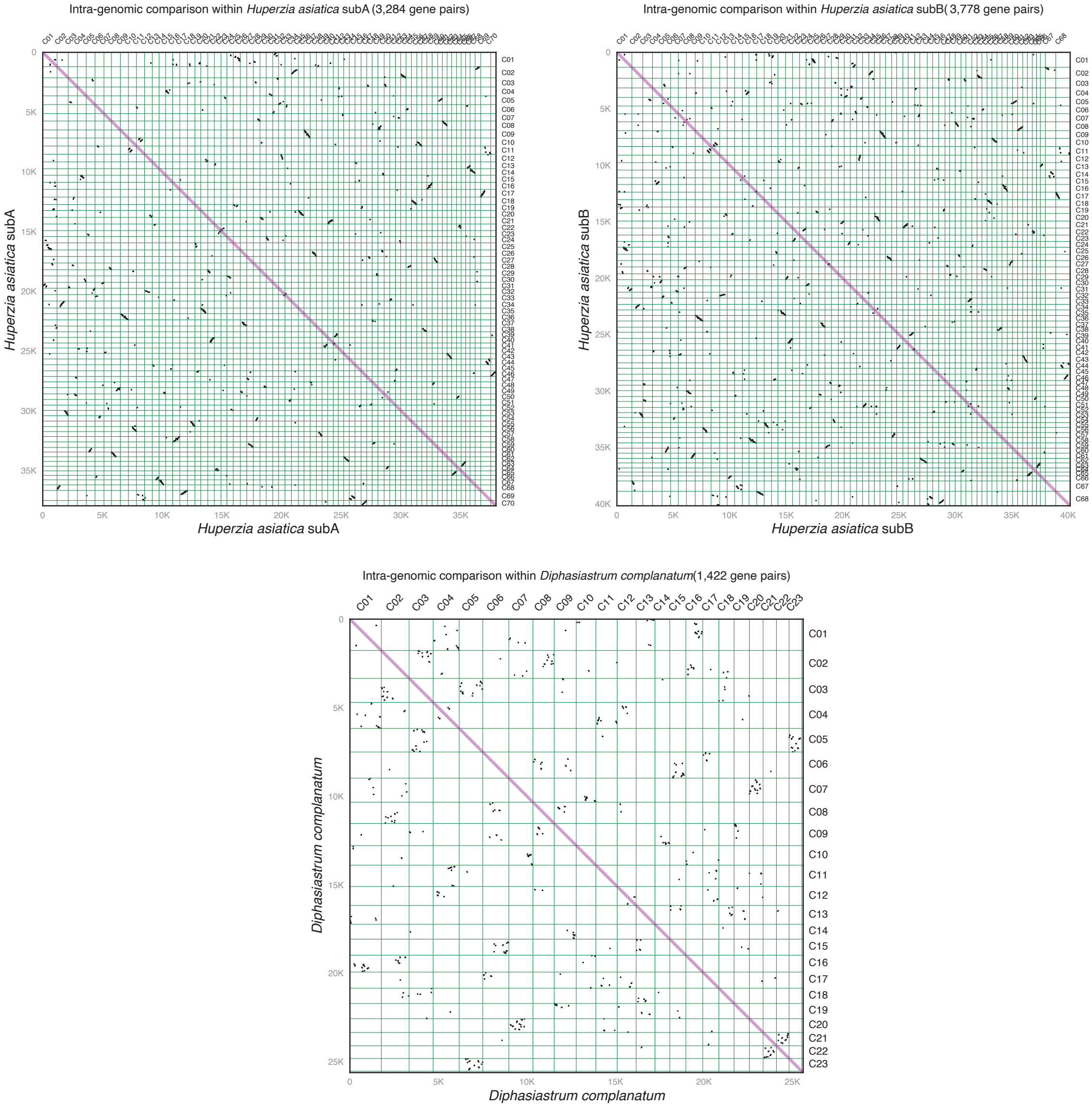


**Supplementary Fig. 13. Plots of intra-genomic synteny in *H. asiatica* and *D. complanatum* genomes.**

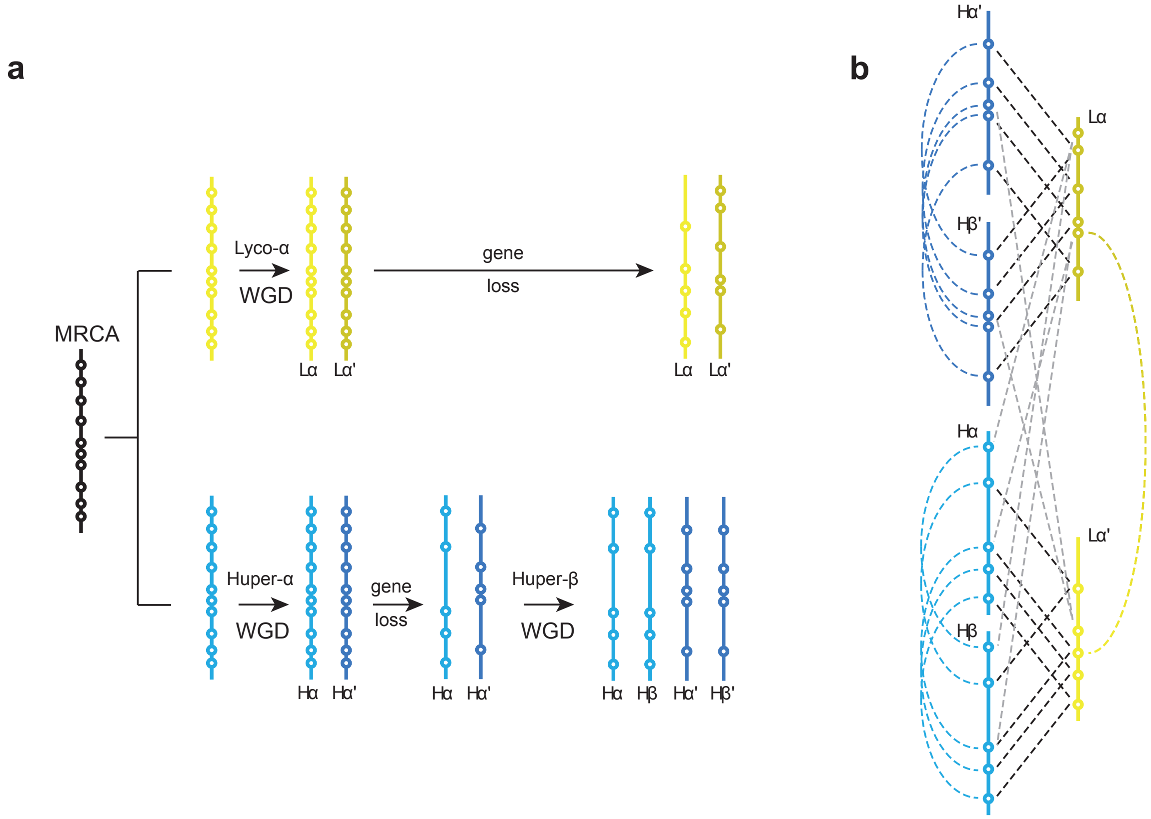


**Supplementary Fig. 14: Diagram of hypothetical syntenic relationships following multiple rounds of WGD and unequal gene fractionation. a**, A hypothetical scenario that could potentially preserve a 2:4 pattern of interspecific synteny between two species while obscuring intraspecific syntenic relationships within them (shown in **b**), similar to what is observed between and within the genomes of *Huperzia asiatica* and *Diphasiastrum complanatum*.

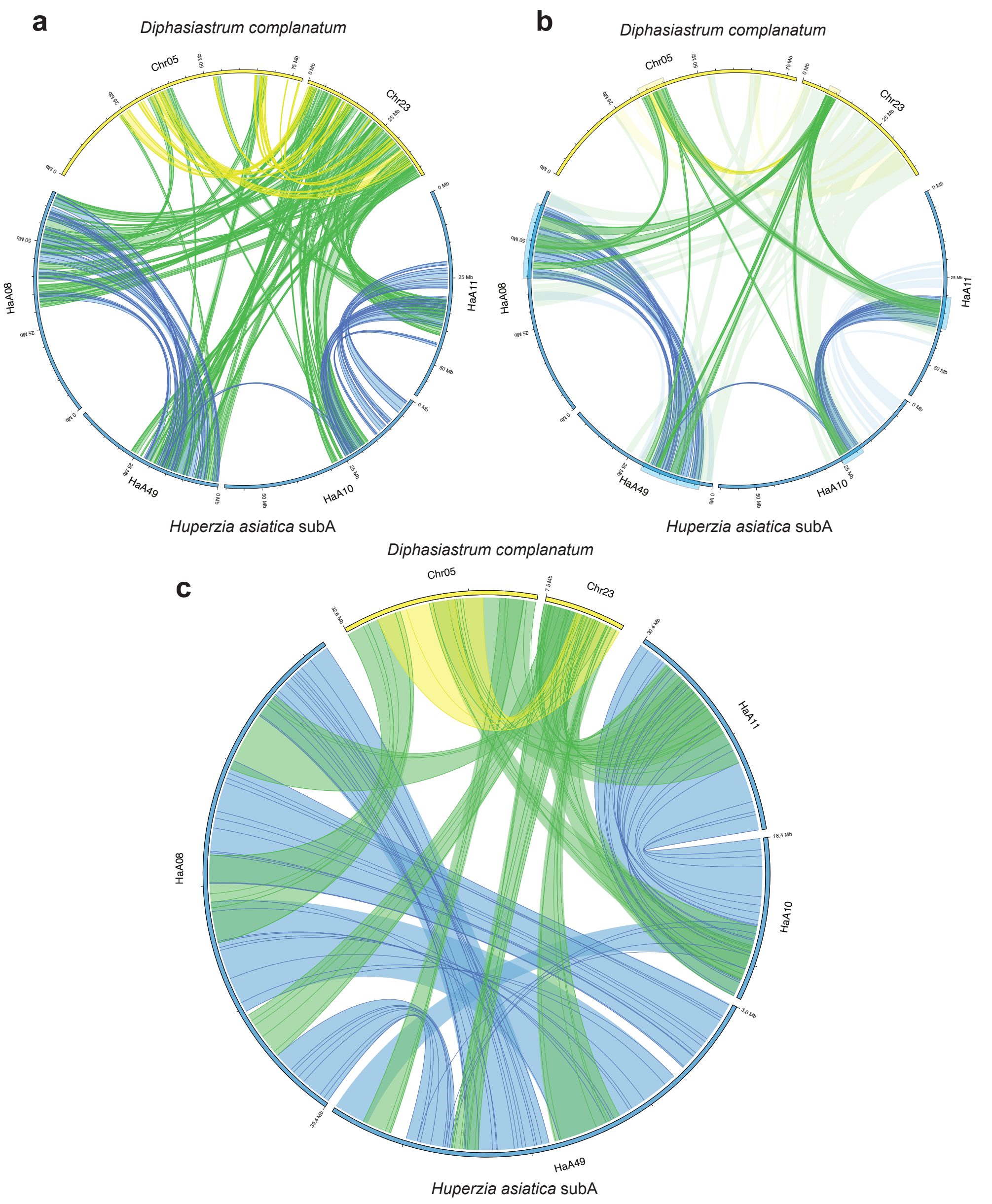


**Supplementary Fig. 15: An in-depth example of collinear blocks exhibiting 2:4 syntenic depth ratio between *Diphasiastrum complanatum* and *Huperzia asiatica* subgenome A.** Circos plots showing: **a,** all syntenic relationships between scaffolds, **b,** a subset of collinear blocks that exhibit a clear 2:4 relationship with regions used for micro-synteny analysis indicated by colored boxes, and **c,** micro-syntenic relationships within the 2:4 region showing overlap of blocks despite little overlap between specific genes within those blocks. Colored ribbons represent blocks of collinear genes with genes represented by darkened lines within each block. Intragenomic relationships are shown in yellow (*D.complanatum*) and blue (*H. asiatica*). Intergenomic relationships are colored green.

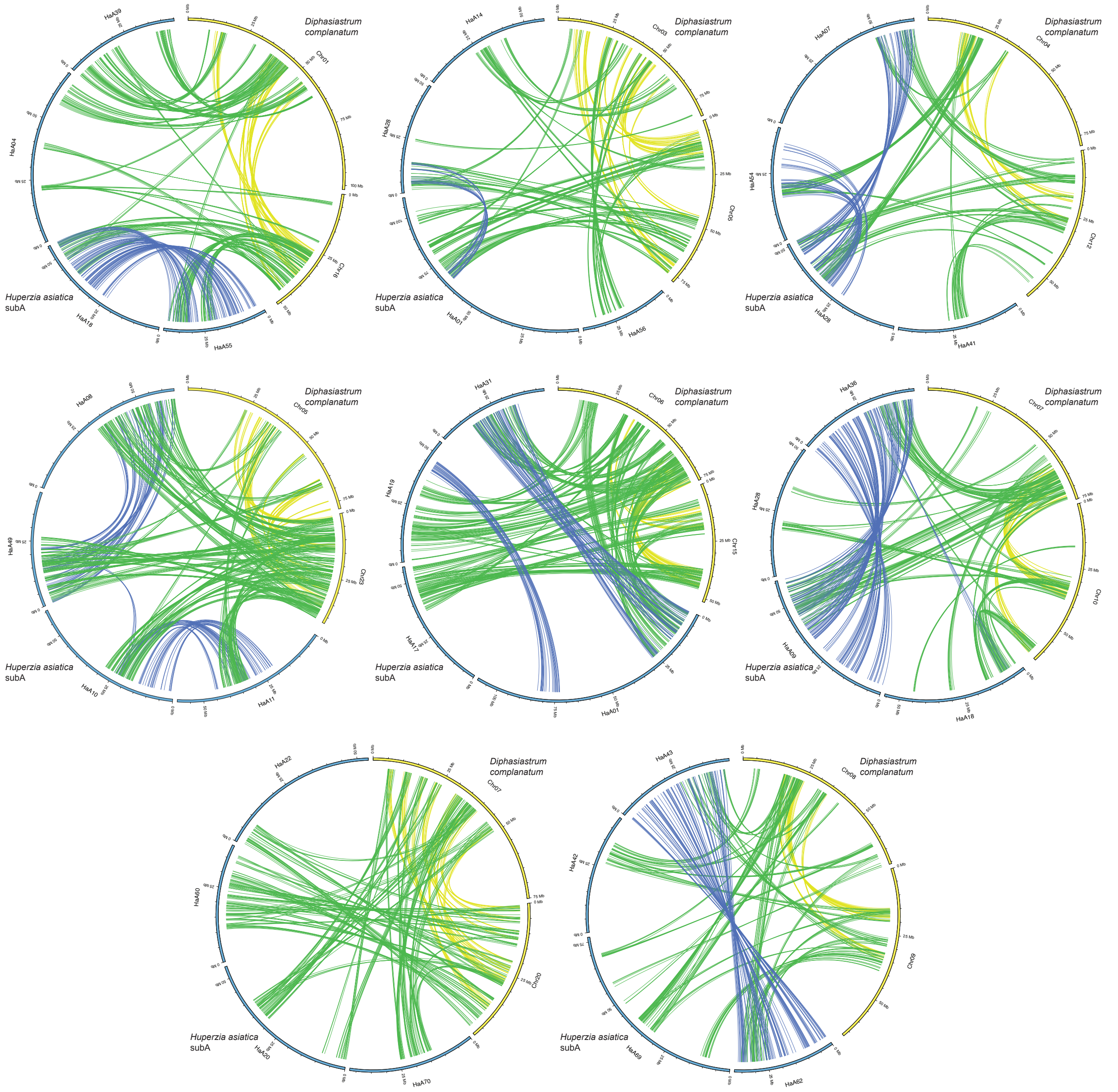


**Supplementary Fig. 16: Collinear genes plotted on scaffolds showing multiple examples of 2:4 syntenic depth between *Diphasiastrum complanatum* and *Huperzia asiatica* subgenome A.** *D. complanatum* scaffolds and intraspecific, collinear links are shown in blue. *H. asiatica* subgenome A scaffolds and intraspecific, collinear links are shown in yellow, and interspecific collinear links are plotted in green.

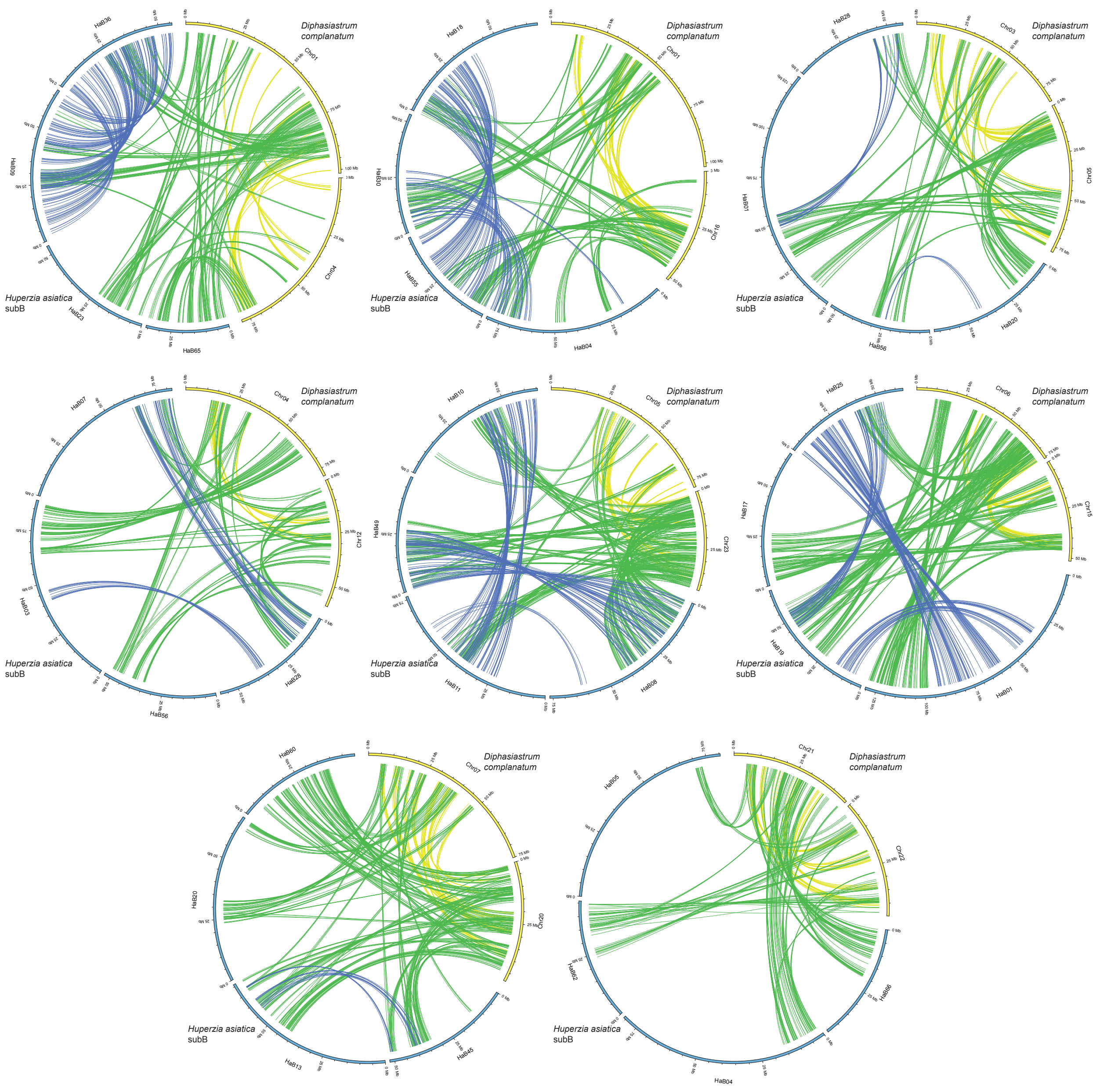


**Supplementary Fig. 17: Collinear genes plotted on scaffolds showing multiple examples of 2:4 syntenic depth between *Diphasiastrum complanatum* and *Huperzia asiatica* subgenome B.** *D. complanatum* scaffolds and intraspecific, collinear links are shown in blue. *H. asiatica* - subgenome B scaffolds and intraspecific, collinear links are shown in yellow, and interspecific collinear links are plotted in green.

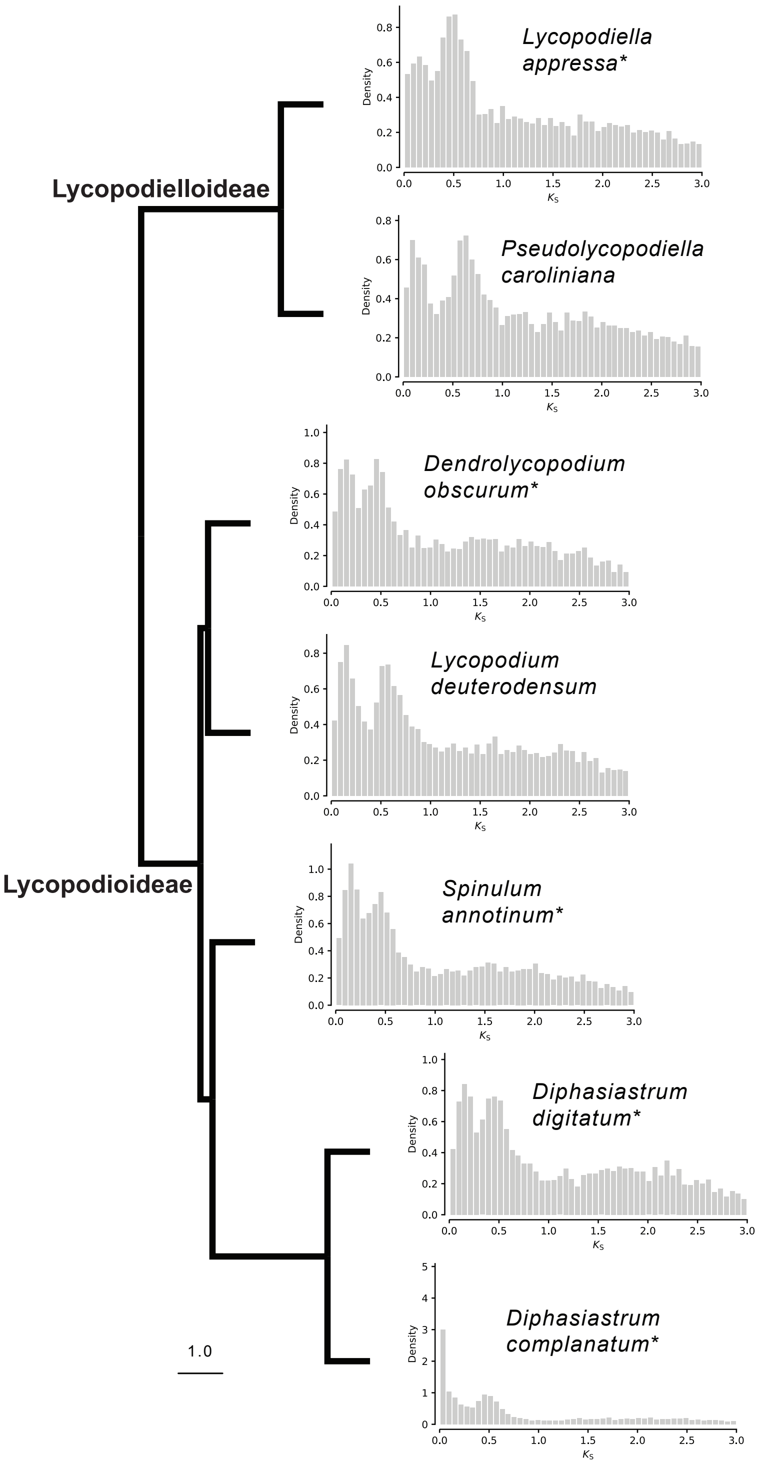


**Supplementary Fig. 18. Ks plots for species in Lycopodioideae and Lycopodielloideae.** Ks plots are displayed on a phylogeny with full ASTRAL support. Species used in MAPs analysis indicated with an asterisk.

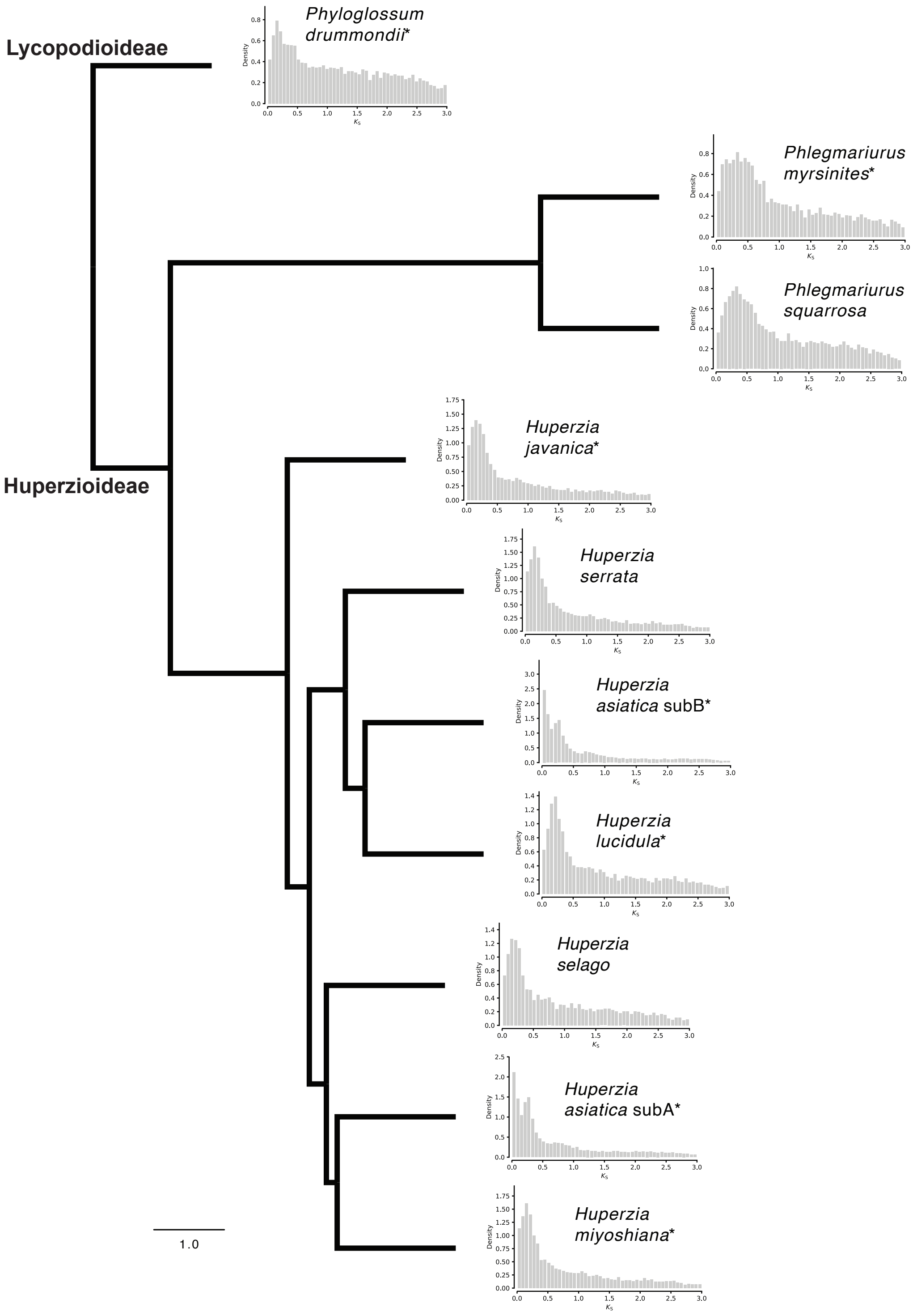


**Supplementary Fig. 19. Ks plots for species in Huperzioideae and Lycopodioideae.** Ks plots are displayed on a phylogeny with full ASTRAL support. Species used in MAPs analyses indicated with an asterisk.

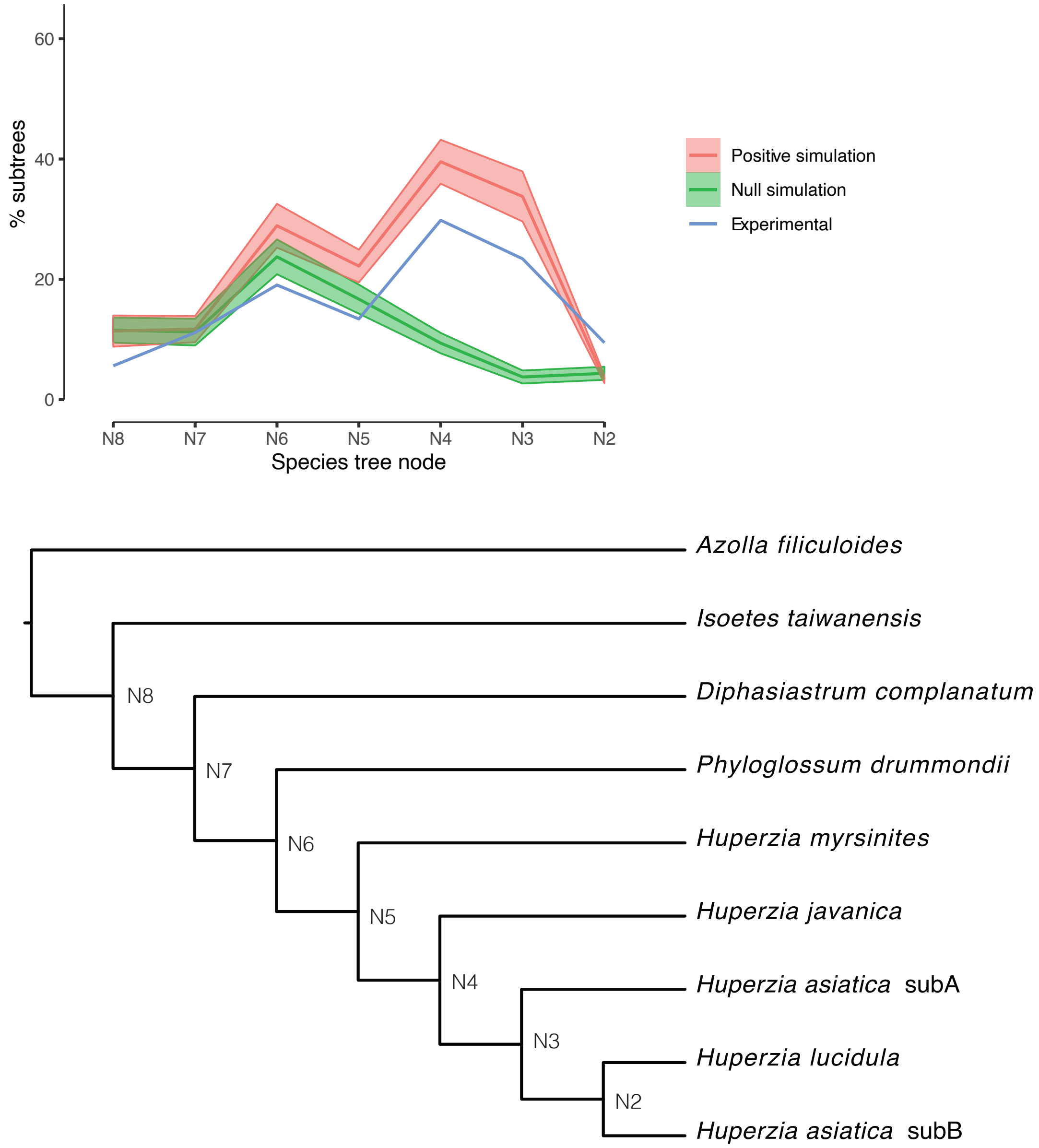


**Supplementary Fig. 20. Summary of MAPs analysis focused on *H. asiatica* subB.** The dark lines in the center of the shaded regions represent the average values for null and positive gene tree simulations. Shaded area shows the standard deviation for gene tree simulations. The taxa sampling here differed from that in Fig. 5d but gave similar results.

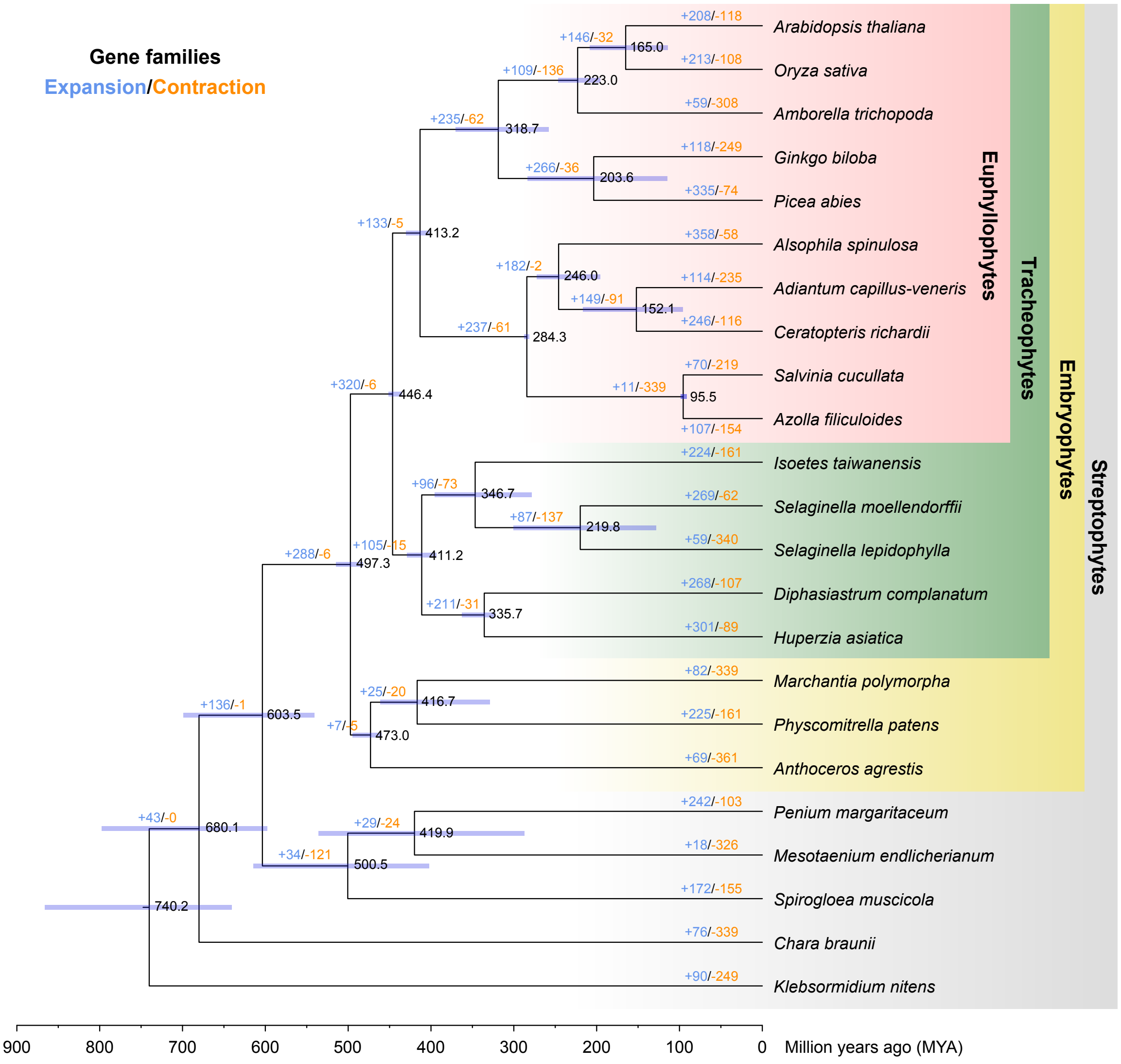


**Supplementary Fig. 21. Gene family evolution of representative land plant lineages.** Numbers in black show the estimated divergence time and 95% confidence intervals at each node in million years ago. Numbers in bule or orange on each branch represent significantly ($P$-value < 0.05) expanded or contracted gene families, respectively, in that branch compared to its last ancestor.

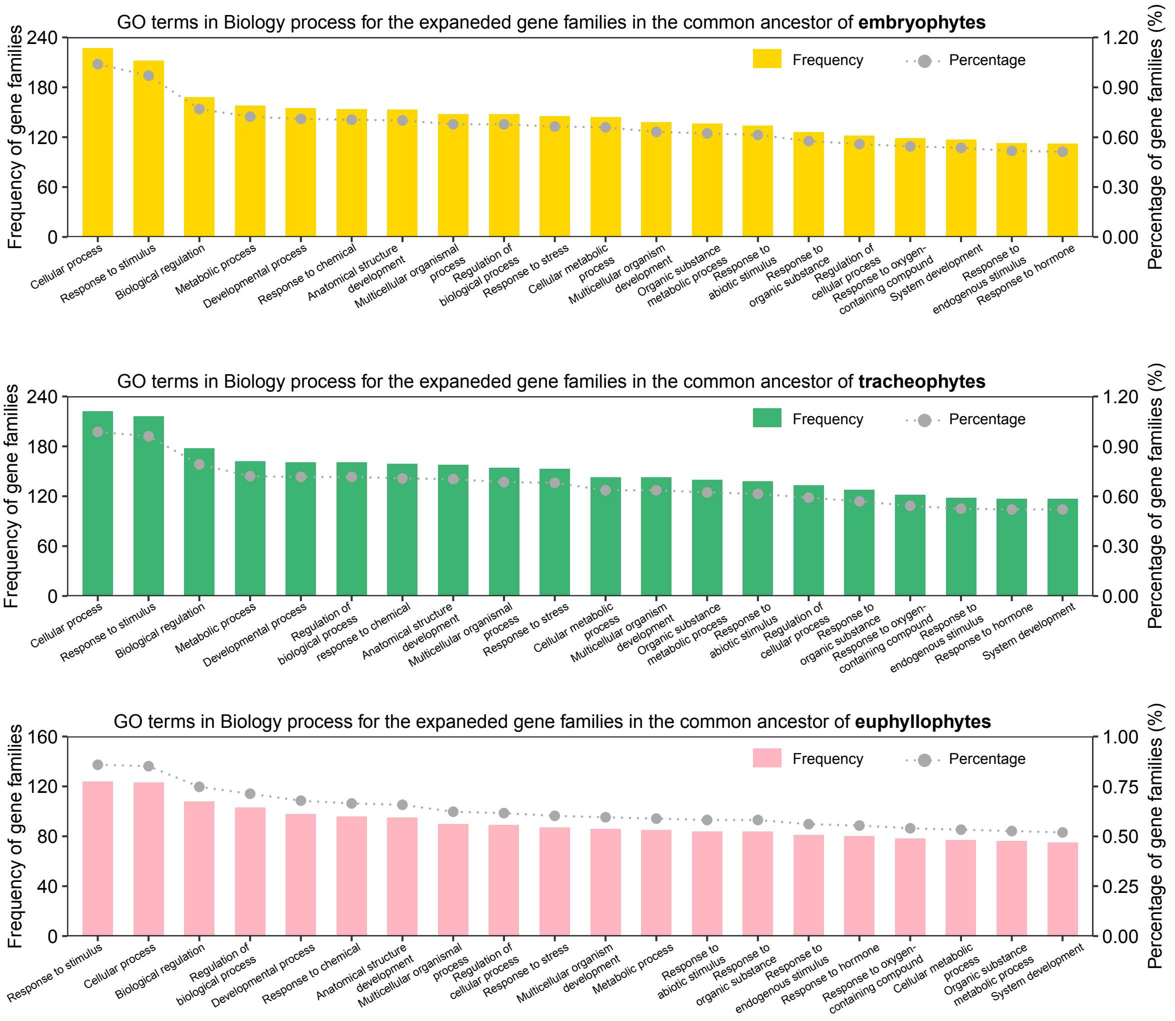


**Supplementary Fig. 22. GO terms in the 'Biological Process' category that are associated with the expanded gene families in the common ancestor of embryophytes, tracheophytes, and euphyllophytes.**

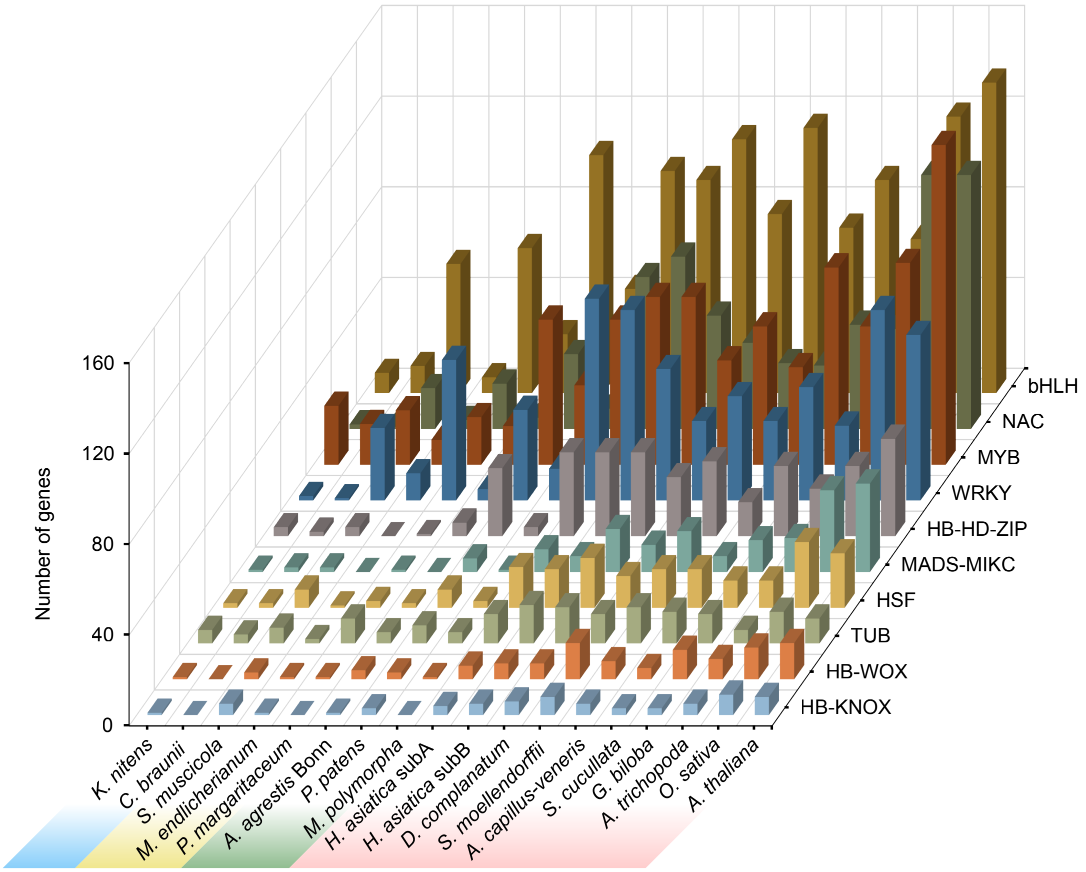


**Supplementary Fig. 23. The copy number of representative transcription factors (TFs) families in land plant lineages.**

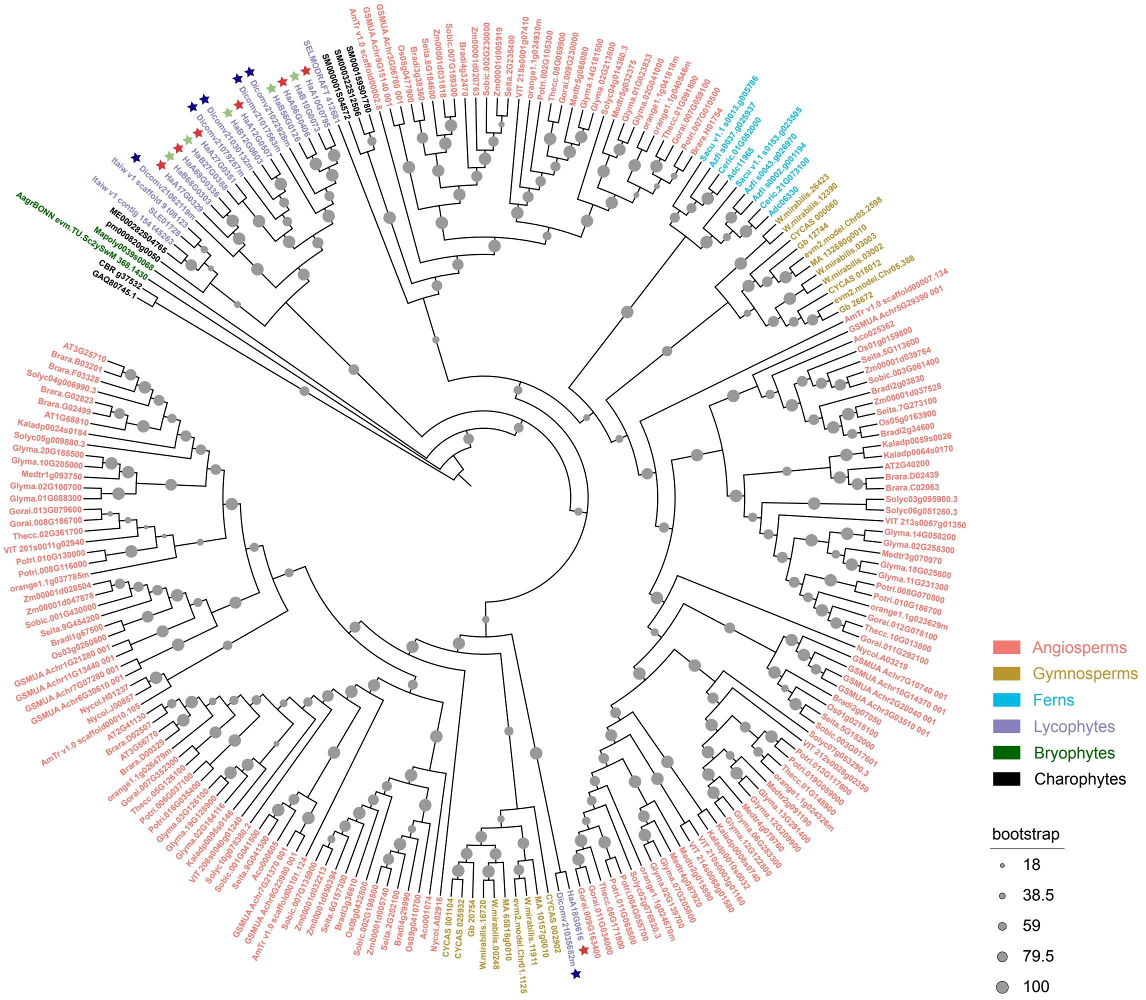


**Supplementary Fig. 24. Phylogenetic analysis of TMO5 proteins in *H. asiatica*, *D. complanatum* and representative species among green plants.** Protein sequences were aligned by MAFFT and the phylogenetic tree was constructed by RAxML using a maximum likelihood method. TMO5 proteins identified in *D. complanatum*, *H. asiatica* subA, and *H. asiatica* subB are marked with dark blue, red, and light green asterisks, respectively.

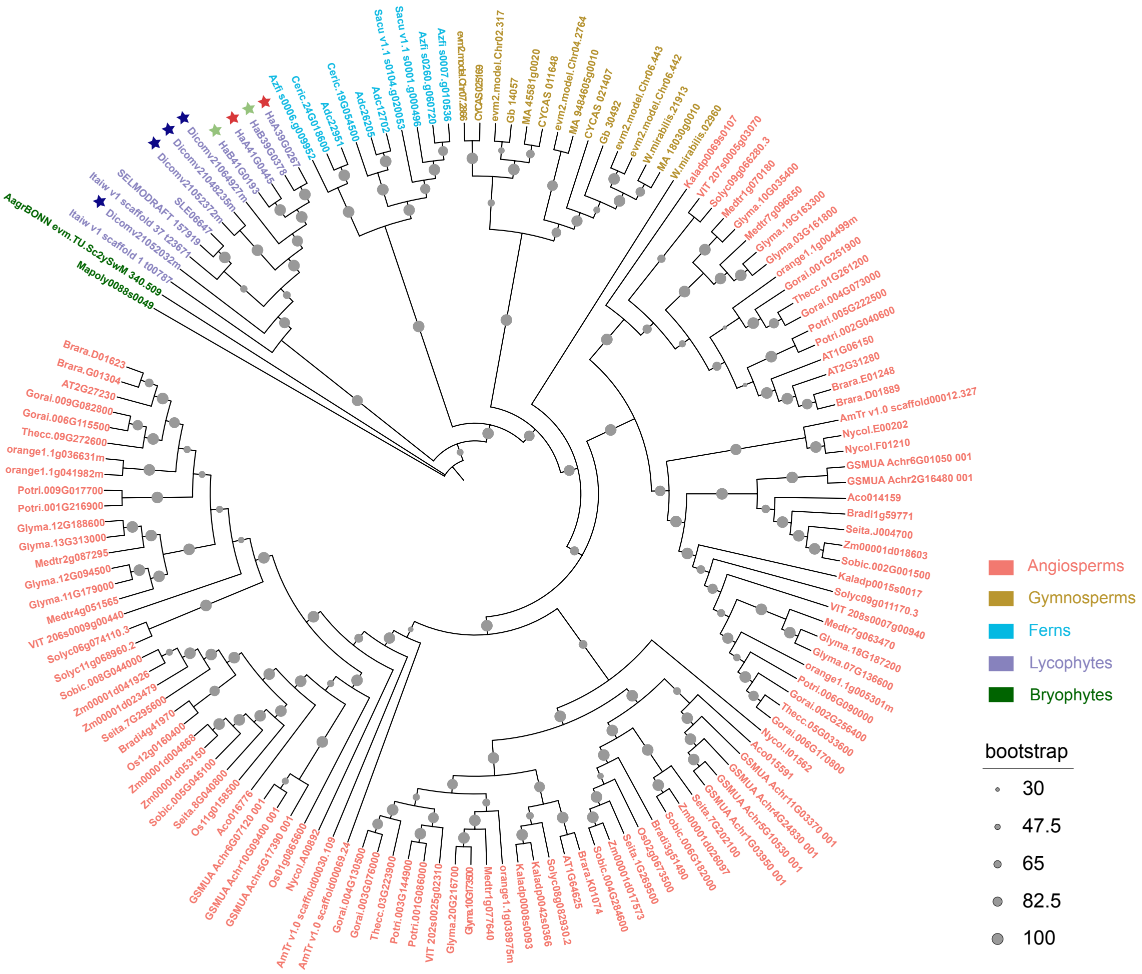


**Supplementary Fig. 25. Phylogenetic analysis of LHW proteins in *H. asiatica*, *D. complanatum* and representative species among green plants.** Protein sequences were aligned by MAFFT and the phylogenetic tree was constructed by RAxML using a maximum likelihood method. LHW proteins identified in *D. complanatum*, *H. asiatica* subA, and *H. asiatica* subB are marked with dark blue, red, and light green asterisks, respectively.

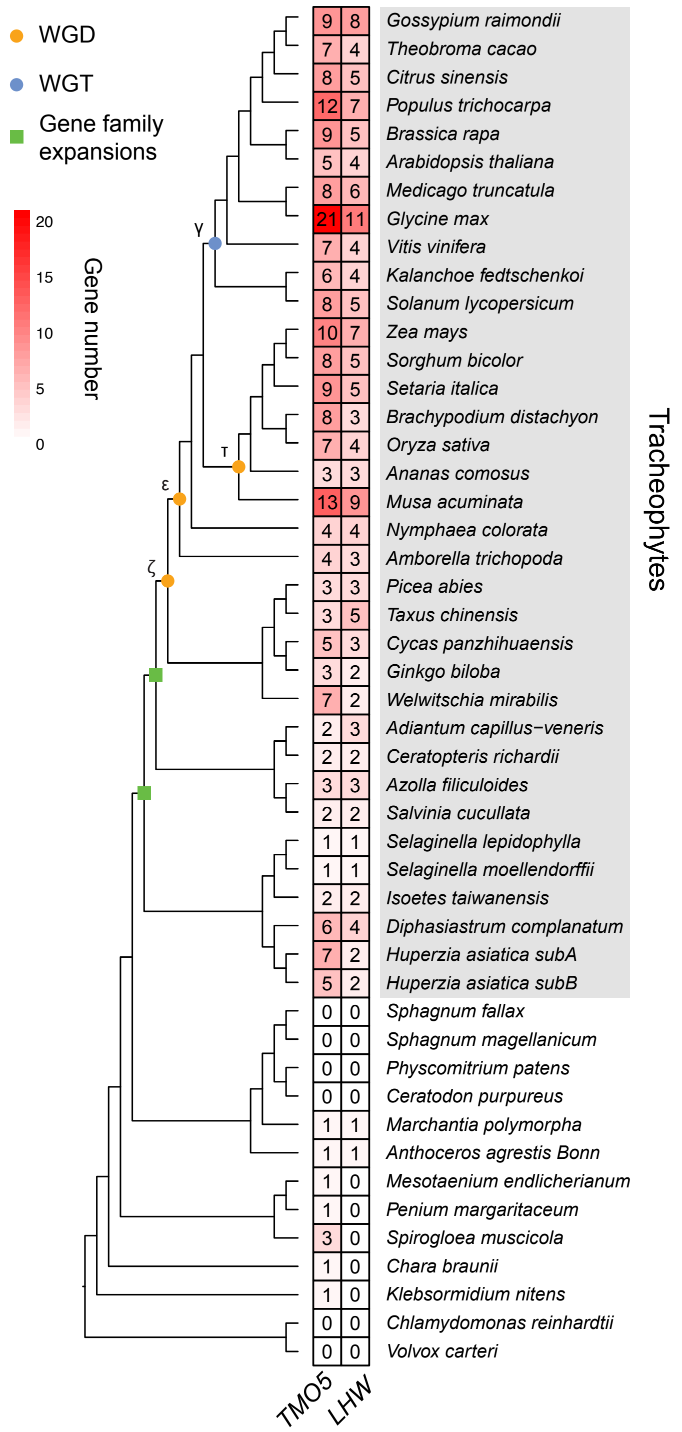


**Supplementary Fig. 26. Phylogenetic profiling of representative species among green plants.** Red-colored cells depict the copy numbers of TMO5 and LHW in each species. WGD, WGT or gene family expansions in the main lineages of vascular plants are marked at each node.
